# Prefrontal activation predicts response latency and is shaped by age and lifestyle

**DOI:** 10.64898/2026.06.22.733821

**Authors:** Rufus Mitchell-Heggs, Daniel Tamkin, Tauheed Ibrahim, Lucas Scherdel, Anita Snowdon-Farrell, Alfred Curry, Onayomi Rosenior-Patten, Simon R Schultz

## Abstract

Neurological and neuropsychiatric conditions affect 43% of the global population, many shaped by modifiable lifestyle exposures, yet their relationship to cortical haemodynamics is poorly characterised. The dorsolateral prefrontal cortex (dlPFC) is a particularly tractable target: it underpins executive function, is disrupted across neuropsychiatric and age-related conditions, and lies on the cortical surface, within reach of scalable optical neuroimaging. We present LUCID, a longitudinal study of 98 healthy adults combining consumer wearable sleep and physical activity metrics with task-evoked dlPFC haemodynamics, measured by time-domain functional near-infrared spectroscopy (TD-fNIRS). Log-transformed peak dlPFC activation was negatively associated with response latency during the Switching Stroop task in both hemispheres (r = −0.32 to −0.38), with stronger associations for the maximum response gradient (r = −0.46 to −0.48), such that greater haemodynamic responses accompanied faster responses. Peak activation showed moderate test–retest reliability (intraclass correlation coefficient, ICC = 0.57–0.69), with between-person variance exceeding within-person fluctuation. Activation was lower in older age groups and higher in females across both tasks. In a subset of 36 participants with complete wearable data, exploratory models combining demographic, lifestyle and scan-quality features predicted bilateral activation under participant-grouped cross-validation (R² = 0.29–0.37), more consistently than task performance, with sex, recent change in REM sleep and age among the strongest participant-level contributors. Together, these findings identify task-evoked dlPFC haemodynamics as a repeatable, behaviourally relevant candidate functional marker, with potential for longitudinal investigation of lifestyle-associated variation in brain function.

## 1 Introduction

Neurological conditions affect 43% of the world’s population (3.4 billion individuals) and are the top contributor to the global disease burden, ahead of cardiovascular diseases (Steinmetz et al., 2024). A growing body of evidence suggests that many of these conditions are not solely the product of genetic predisposition or acute injury, but emerge from the cumulative effects of modifiable lifestyle factors over time.

A common thread linking many of these conditions is disruption to cerebral haemodynamics, specifically the delivery and regulation of oxygenated blood flow to the brain. Such disruption is observed across vascular dementia, traumatic brain injury (TBI), ischaemic stroke, schizophrenia, and neurodegenerative diseases, where alterations in the haemodynamic response serve as both a marker of underlying pathology and a potential target for monitoring disease progression or recovery (Patel, Nischal and Patel, 2026). This pattern extends to neuropsychiatric conditions: systematic review and meta-analysis evidence indicates that depressed patients show a consistent reduction in cerebral blood flow (CBF) relative to healthy controls, suggesting a role for haemodynamic assessment and CBF-altering interventions in high-risk groups (Chithiramohan et al., 2022). In attention deficit hyperactivity disorder (ADHD), prefrontal hypoactivation during executive function tasks is consistently observed using functional near-infrared spectroscopy (fNIRS), with diminished oxyhaemoglobin (HbO) responses in the prefrontal cortex (PFC) documented in both child and adult populations (Peet et al., 2022; Zhang et al., 2023), consistent with the broader pattern of haemodynamic disruption observed across neurological and neuropsychiatric conditions.

Critically, however, many of these conditions do not arise acutely. The concept of the exposome, the totality of environmental and lifestyle exposures an individual accumulates across their lifespan offers a framework for understanding this incremental trajectory, capturing how cumulative exposures interact with genetic background to shape brain health and disease vulnerability over time (Sakowski et al., 2024; *Nature Reviews Neuroscience*, 2026). In dementia, for example, the 2024 Lancet Commission identified fourteen modifiable risk factors spanning physical, social, and environmental domains that, if addressed from childhood through later life, could prevent or delay nearly half of all cases worldwide (Livingston et al., 2024). This principle extends across neurological conditions more broadly: for a substantial proportion of cases, the trajectory toward disease is shaped by compounding everyday lifestyle exposures rather than discrete clinical events.

Haemodynamic imaging has predominantly been applied in the context of established diagnosis, yet the more pressing opportunity lies upstream - identifying and continuously tracking the lifestyle factors that cumulatively modulate cerebral haemodynamics before clinical thresholds are reached. The dorsolateral Prefrontal Cortex (dlPFC) is a particularly compelling target. It underpins the core executive functions, working memory, cognitive flexibility, planning, and inhibitory control (Jung et al., 2022) - which are themselves sensitive to lifestyle-related variation in sleep and physical activity (Csipo et al., 2021; Shen et al., 2024). Its haemodynamic function is disrupted across neuropsychiatric and age-related conditions (Chithiramohan et al., 2022; Peet et al., 2022; Zhang et al., 2023; Patel, Nischal and Patel, 2026) and, as a superficial cortical region, it is also accessible to reliable, scalable optical neuroimaging (Ban et al., 2022; Castillo et al., 2024; Taylor et al., 2024).

Recent work has begun to address this opportunity, establishing that sleep and physical activity each exert measurable effects on cortical haemodynamics. Sleep deprivation has been shown to significantly alter task-evoked prefrontal haemodynamic responses and reduce task-associated cerebral blood flow, with concurrent impairments in response latency (RT) and sustained attention (Csipo et al., 2021; Rab-Bábel et al., 2025). Physical activity interventions produce acute increases in prefrontal oxyhaemoglobin, with effects varying by exercise type, intensity, and duration (Naito et al., 2024), and are associated with improvements in working memory and inhibitory control (Shen et al., 2024). Where both domains have been examined together, their contributions to brain function appear distinct: sleep and physical activity relate specifically to brain connectivity during cognitively demanding tasks, while heart rate variability and respiration rate are more relevant for resting-state connectivity (Triana et al., 2024).

However, two critical limitations constrain the translational value of this literature. First, most are single-session or acutely manipulated designs. These capture transient state effects rather than the stable individual differences that arise from habitual lifestyle patterns, making it impossible to separate trait-level haemodynamic sensitivity from day-to-day fluctuation. Second, prior work has rarely targeted the dlPFC specifically: prefrontal effects have typically been captured with broad cortical coverage or framed around connectivity, rather than as targeted, task-evoked activation under standardised executive load.

The present study addresses both limitations with a longitudinal, repeated-measures design spanning three sessions in a healthy adult cohort. We pair task-evoked dlPFC haemodynamics, measured by time-domain functional Near-infrared Spectroscopy (TD-fNIRS), with quantitative, multimodal lifestyle data from consumer wearables. Consumer-grade wearable devices now make it feasible to continuously capture a broad suite of lifestyle-relevant metrics longitudinally and at scale, providing objective proxies for the cumulative lifestyle exposures that may shape prefrontal brain function. This study focuses specifically on sleep and physical activity, two of the most consistently implicated modifiable lifestyle domains captured quantitatively across multiple temporal windows prior to each scanning session. Complementing this, TD-fNIRS enables direct measurement of dlPFC oxygenation in a portable, repeated-measures protocol that is well-suited to longitudinal multimodal study designs. Critically, TD-fNIRS is considered the gold standard of non-invasive optical brain imaging, offering superior depth sensitivity and improved quantitative estimates of oxyhaemoglobin concentration relative to conventional continuous-wave fNIRS systems, providing greater confidence that measured haemodynamic signals reflect cortical rather than superficial tissue changes (Ban et al., 2022; Castillo et al., 2024).

Using this framework, we continuously and quantitatively captured sleep and physical activity metrics alongside task-evoked dlPFC haemodynamics and standardised cognitive assessments, the Switching Stroop task and the 2N-Back working memory test, across three repeated sessions in a healthy adult cohort. Unlike prior work relying on self-report or single-domain monitoring, this design enables characterisation of stable individual differences in prefrontal brain state and their relationship to habitual lifestyle behaviour. We report three principal findings. First, peak dlPFC oxyhaemoglobin activation, measured via TD-fNIRS, was associated with response latency during the Switching Stroop task, with greater activation accompanying faster responses. Across both executive tasks, activation showed moderate test-retest reliability, with between-person variance exceeding within-person fluctuation. Second, activation varied with age and sex across the full cohort, with lower activation in older participants and higher activation in females. In an exploratory subset of 36 participants with complete wearable data, recent changes in REM sleep duration also contributed to activation prediction alongside demographic and scan-quality features. Third, within this subset, these features predicted haemodynamic activation more consistently than task performance, while composites combining haemodynamic response and behaviour offered no improvement over activation alone within the tested models. Together, these findings identify task-evoked dlPFC activation as a repeatable, behaviourally relevant candidate functional marker for longitudinal investigation of lifestyle-associated variation in brain function.

## 2 Methods

### 2.1 Participants

Ninety-eight healthy adults (47 female; age range 18–55 years) were recruited via mailing lists, social media, and local institutional networks. Eligibility was assessed through an anonymous online pre-screening questionnaire (Qualtrics). Eligible respondents were contacted by email, provided with a Participant Information Sheet before booking their first appointment, and gave written informed consent at the first session after confirming they had read and understood the study information.

Eligibility criteria were: aged 18–55 years; normal or corrected-to-normal vision and hearing; fluency in English; at least one month of consistent consumer wearable use (e.g. Oura, Whoop, Apple Watch, Garmin, or Fitbit), with consent to share data before and during the study; availability to attend up to three imaging sessions at Imperial College London; and willingness to comply with study procedures, including wearing the Kernel Flow 2 TD-fNIRS headset and completing the cognitive task battery.

**Table 1:**
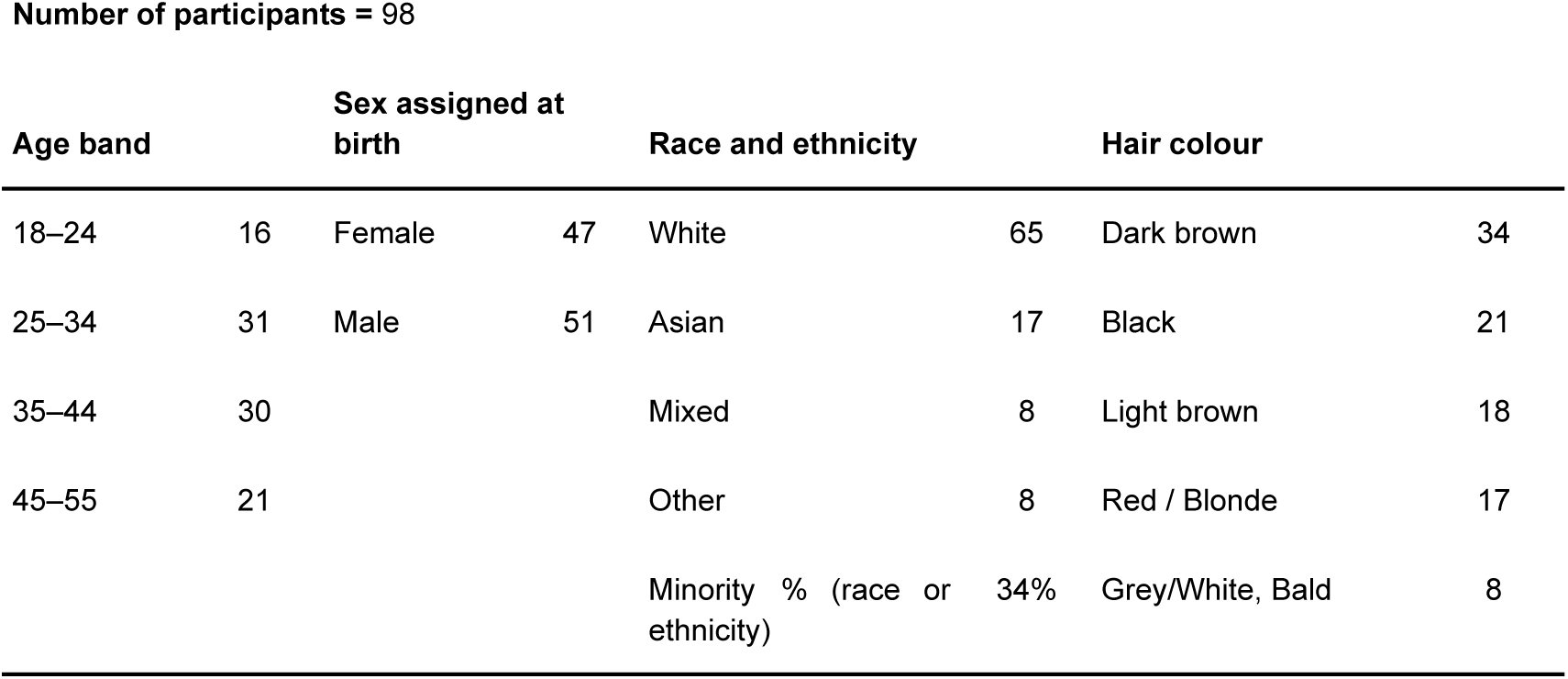
Demographic characteristics of the LUCID cohort (N = 98), summarising age band, sex assigned at birth, race and ethnicity, and hair colour.

Exclusion criteria were: history of neurological conditions (including brain surgery, craniotomy, epilepsy, stroke, or concussion with loss of consciousness); history of acute migraine; active psychiatric condition; major comorbid or unstable medical condition; current use of psychoactive medication; pregnancy (self-reported); and inability to provide informed consent or follow task instructions due to severe cognitive or language impairment.

### 2.2 Ethical approvals

This study received favourable opinion from the Imperial College Research Ethics Committee (ICREC; ref. 7657797) and departmental approval from the Head of the Department of Bioengineering. All researchers involved in the study hold Good Clinical Practice (GCP) certification, in compliance with the UK Policy for Health and Social Care Research.

### 2.3 Wearable and lifestyle features

#### 2.3.1 Wearable data Acquisition

Lifestyle and physiological data were collected using wearable devices integrated via the Terra API. Data streams were available for up to three months prior to the first scan and throughout the study period.

Collected variables included:

1. Physical activity (e.g., steps, energy expenditure)
2. Sleep (duration, efficiency, latency, regularity)
3. Cardiovascular measures (resting heart rate, heart rate variability)
4. Respiratory and oxygenation metrics

**Table 2.** N=36 Participants selected for further wearable analysis due to data availability.

| Wearable breakdown |  |  |  |  |
| --- | --- | --- | --- | --- |
| Whoop | Apple Watch | Oura | Fitbit | Garmin |
| 10 | 1 | 7 | 1 | 17 |

#### 2.3.2 Wearable metric extraction

Daily-resolution wearable metrics spanning activity, sleep, and vital-sign domains were retrieved for each participant, using only records that ended within the 90 days preceding each scan. Non-numeric metrics (for example, sleep end time) and sleep sessions flagged as naps were excluded. Each sleep record was dated by the morning it ended, and each daily record by the day it covered. Where a participant had data from more than one provider, a single provider was used for each scan, chosen as the one contributing the most records in the window, with Apple used only when no other provider had data. Where a metric had more than one record on the same day, the longest record was kept.

Wearable metrics were filtered on coverage at two levels. Within each participant, a metric was used for a given scan only if it had been recorded on at least 30 of the 90 days before the scan, so that a baseline could be calculated. Across the cohort, a metric was retained only if at least 40 participants met this criterion before their first scan (Supplementary Material A.1).

#### 2.3.3 Calculation of wearable features

For each surviving metric, we derived trait features, describing the participant’s habitual level, trend and variability across the 90 days before the scan, and state features, describing their departure from these immediately before the scan. Separating each metric into trait and state components follows the disaggregation of between-person and within-person effects recommended for repeated-measures data (Curran and Bauer, 2011). Most features were scaled by the participant’s typical spread, s, defined as 1.4826 × the median absolute deviation from the median of their data (the scaling factor approximates the standard deviation), so that features are expressed relative to each participant’s own variability.

Four trait features were computed. The baseline was an exponentially weighted moving average with a 30-day half-life, with weights decaying with elapsed time rather than number of observations, evaluated at the most recent day; this weights recent days more heavily, so that the baseline reflects the participant’s current habitual level rather than an equal average over the full window. The slope captured whether the metric was trending up or down, estimated with the Theil-Sen estimator, a robust linear slope that is resistant to outliers, and divided by s, with positive values indicating an upward trend. The coefficient of variation (CV) was s divided by the median. The root mean square of successive differences (RMSSD) was computed from the differences between values on consecutive days, ignoring pairs separated by a missing day, and divided by s.

Three types of state features were computed. The day deviation was the value on the night before the scan (sleep metrics) or the last full day before the scan (daily metrics), minus the baseline, divided by s. The moving average convergence divergence (MACD) was the difference between exponentially weighted moving averages with 7-day and 30-day half-lives, divided by s, with positive values indicating that the recent level was above the long-term level. Finally, recent-change features were computed for slope, RMSSD and CV as the same statistic over the last 10 days before the scan minus its value over the full 90 days, with the slope and RMSSD differences also divided by s. MACD and the recent-change features additionally required at least 3 observations of the metric in the last 10 days before the scan.

#### 2.3.4 Coverage of wearable features across providers

Since we collected data from multiple provider brands, we aimed to reduce systematic biases between devices. We removed metrics that were not reported by all four of the most common providers in our sample (Apple, Garmin, Oura and Whoop), as well as those with known reporting differences between providers. Sleep heart rate variability was retained although Apple reports SDNN whereas the other providers report RMSSD; we treated the two as equivalent and used RMSSD wherever it was available. We then compared the baseline and CV of each remaining metric across providers using Kruskal-Wallis H tests, and excluded any metric for which both features differed between providers at p < 0.001 (Supplementary Material A.2).

#### 2.3.5 Clustering of wearable metrics

Different wearable metrics are naturally correlated with each other (e.g., activity and heart rate), which would introduce redundancies in models of the impact of the wearables on dlPFC activity. We therefore used hierarchical clustering to group correlated metrics together, so that at most one metric from each cluster could enter the models (see 2.5.6).

To build the clusters, we first estimated how strongly each pair of metrics co-varied within participants, using the same daily values as feature extraction (2.3.2). For each participant with at least 30 days of data, we calculated the Spearman correlation between each pair of metrics across the days on which both were recorded, requiring at least 30 shared days. Correlations were Fisher z-transformed, averaged across participants weighted by n - 3 (where n is the number of shared days), and transformed back. Metrics were then clustered by average linkage on the distance 1 - |r|, and the tree was cut into three clusters (Supplementary Material A.4).

### 2.4 Cognitive Assessment Protocol

Each participant completed three sessions separated by approximately 7–10 days. Each session followed a standardised protocol comprising: a pre-scan contextual survey; a cognitive task battery, and a post-scan survey. Cognitive performance was assessed using a battery of tasks implemented via Cognitron (formerly H2 Cognitive Designs; Hampshire et al., 2019). Participants completed a 4-minute eyes-closed rest period prior to the task battery. Tasks were then administered in a fixed order: Immediate Object Recall, Digit Span, 2N-back, Stroop, and Delayed Object Recall with a 15 second eyes-open rest period between consecutive tasks. A second 4-minute eyes-closed rest period was completed following the task battery. A subset of which was used for the present analysis (2N-Back and Switching Stroop (detailed below).

#### 2.4.1 2N-Back task paradigm

Working memory was assessed with a 2N-Back paradigm (see Figure 2a, left). Participants viewed a sequential stream of images (N = 78 trials) and indicated, for each image, whether it matched the image presented two positions earlier. Images were drawn from a stimulus set and responses were made via response mapping “Match”/“No Match” button press. A different image sequence was used in each session. RT (from stimulus onset to response) and match accuracy were recorded as outcome measures. All trials were aggregated for task-evoked haemodynamics.

**Figure 1.**
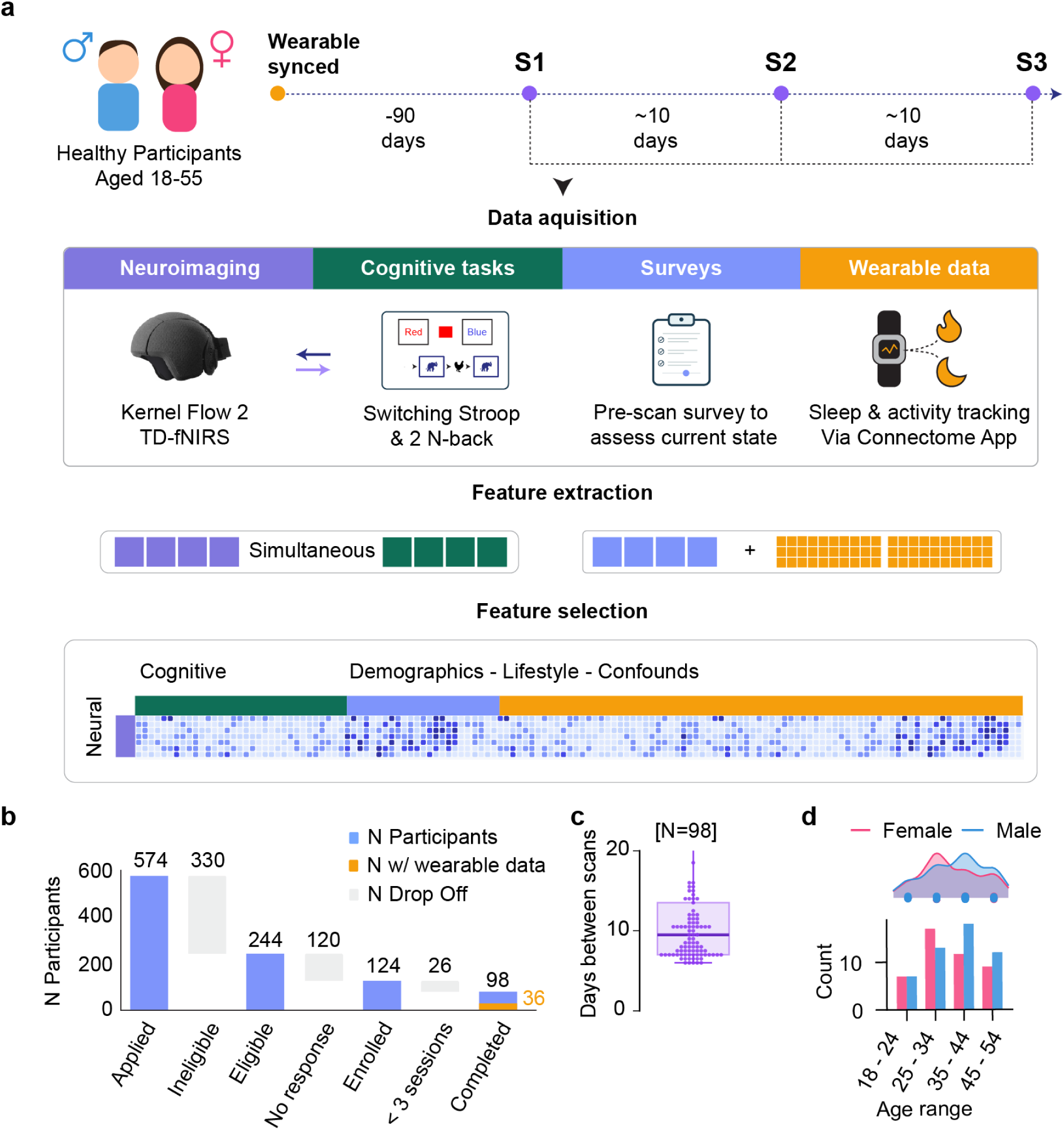
Study design, data acquisition and feature extraction pipeline. **(a)** Schematic overview of the longitudinal study protocol in healthy participants aged 18–55. Wearable devices were synced approximately 90 days prior to the first session to establish baseline data. Participants completed three in-person sessions (S1–S3), each separated by approximately 10 days. At each session, four complementary data modalities were integrated: (i) neuroimaging using the Kernel Flow 2 TD-fNIRS system; (ii) cognitive performance from a Switching Stroop task and a 2N-Back working memory task; (iii) surveys capturing self-reported contextual state; and (iv) longitudinal wearable-derived sleep, physical activity, and physiological metrics collected before and throughout the study period. Following data acquisition, neural and cognitive features were extracted simultaneously from each session. Feature selection was then applied to identify the most informative variables across cognitive, neural, and composite (demographic, lifestyle, and confound) feature sets for downstream modelling. **(b)** Participant eligibility flow: 574 applicants, 330 were ineligible and 120 of the remaining 244 did not respond, giving 124 enrolled; 26 completed fewer than three sessions, yielding a final sample of 98 for dlPFC marker analysis (figures 2, 3), of whom 36 (108 scans) had complete wearable data and formed the ridge model analysis (figure 4). **(c)** Distribution of mean inter-session intervals (in days) across the final sample (N = 98). The box plot shows the median, interquartile range, and full spread of session spacing, with individual participant values overlaid as scatter points. **(d)** Age and sex distribution of the final sample. Participants are grouped into ten-year age bands (18–24, 25–34, 35–44, 45–55) and counts are shown separately for female (pink) and male (blue) participants, with a density overlay illustrating the overall distribution by sex.

**Figure 2.**
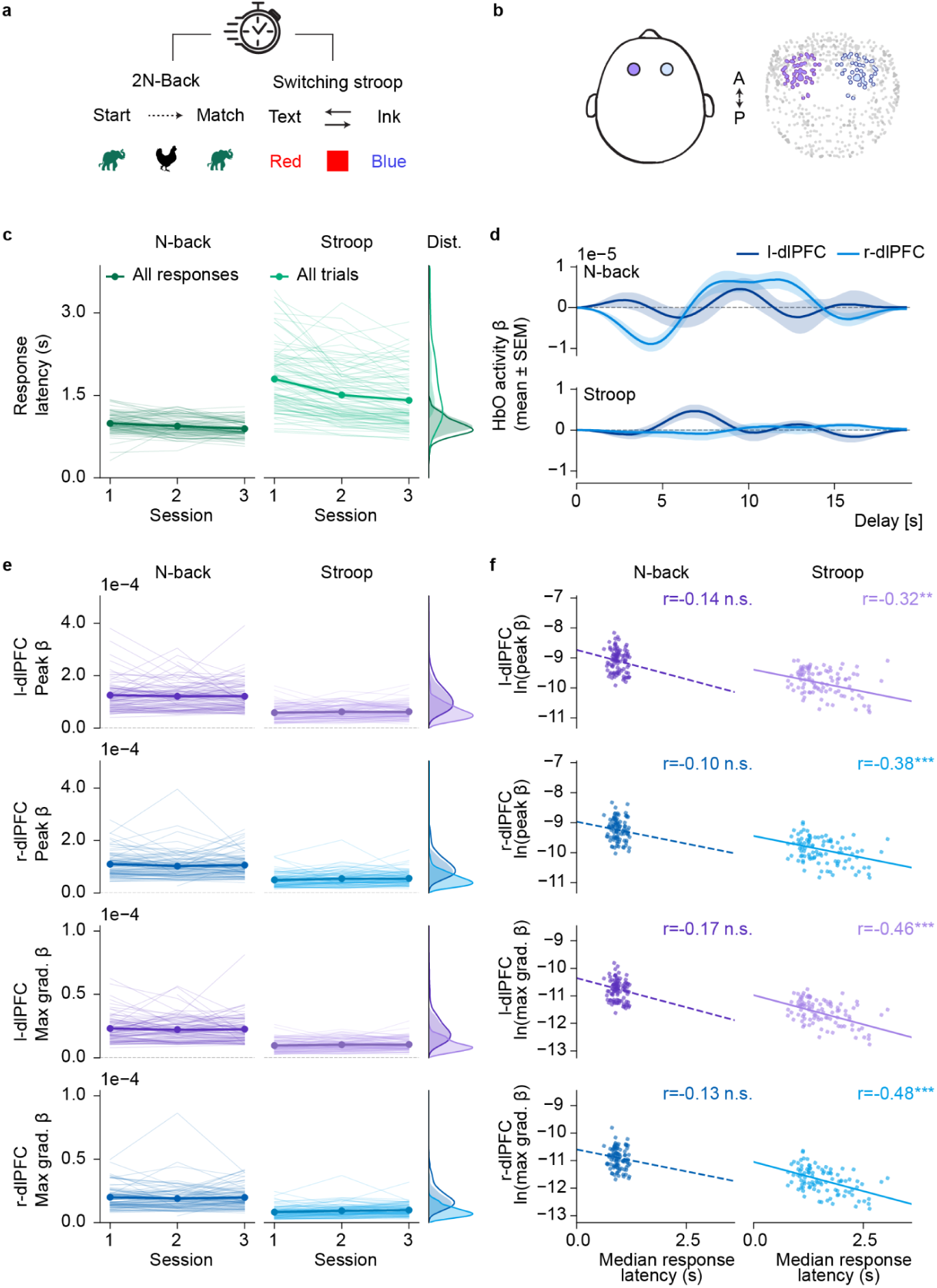
Greater dlPFC haemodynamic activity is linked to faster Stroop response latency. (a) Task paradigms. 2N-Back (left), participants are presented with one image at a time and asked to select Match or No Match. Switching Stroop (right), participants are presented with one of two rules and select the option (Red or Blue) that corresponds to the rule. (b) Simultaneous TD-fNIRS of left and right dlPFC whilst completing the tests. (c) Mean and individual median response latency per session, measured from stimulus presentation to decision click, for 2N-Back (left; all responses) and Stroop (right; all trials), with the distribution across sessions. (d) Group mean ± SEM of the task-evoked HbO response (FIR β) for left and right dlPFC. (e) Peak β and maximum gradient β across sessions (S1 to S3), for each hemisphere and task. Thin lines are participants and bold markers are session means. (f) Log peak β and log maximum gradient β against median response latency, one point per participant mean. Lines show linear fits, solid when p < 0.05 and dashed otherwise. Stroop associations were significant for peak β (left r = −0.32, right r = −0.38) and maximum gradient β (left r = −0.46, right r = −0.48), all p < 0.01, whereas 2N-Back associations were not (r = −0.10 to −0.17).

#### 2.4.2 Switching Stroop task paradigm

Participants completed a Switching Stroop colour-word interference task (N = 60 trials; Figure 2a, right). The task incorporated two response rules, requiring participants to respond according to either the written word or its ink colour, as indicated by a task cue. Participants selected “Red” or “Blue” by clicking the corresponding response box. Response latency (RT) and accuracy were recorded for each trial type, and condition contrasts (incongruent minus congruent; switch minus non-switch) were computed. All trials were aggregated for the task-evoked haemodynamic analysis.

Only the 2N-Back and Switching Stroop tasks were included in the current analysis (see methods for task-evoked haemodynamic analysis). An important note, is that these tasks differed in their response latency demands: the 2N-Back paradigm imposed external time pressure (i.e. the task stimulus window would move on if the participant didn’t select anything), whereas Stroop RTs reflect self-paced, deliberate response selection (i.e. the participant had to click an option for the test to continue).

### 2.5 TD-fNIRS feature extraction

#### 2.5.1 TD-fNIRS Data Acquisition

Neuroimaging data were acquired with the Kernel Flow2, a second-generation TD-fNIRS system (Kernel, Culver City, CA, USA) (Ban et al, 2022). At the start of each session, the headset was calibrated and positioned so that the anterior edge of the frontal modules aligned with the top of the participant’s eyebrows, ensuring consistent placement over the dlPFC across sessions. Unlike continuous-wave fNIRS, which records only the attenuation of light, TD-fNIRS emits picosecond light pulses and measures the full distribution of photon times-of-flight (DTOF) at each channel. Because late-arriving photons preferentially sample deeper tissue, resolving this temporal profile affords greater depth sensitivity and a partial separation of superficial extracerebral signal from cortical signal. The system illuminated tissue at two wavelengths (690 and 905 nm) and recorded at an effective sampling rate of 3.76 or 4.76 Hz (the acquisition rate was changed by the manufacturer during the study period). Each module comprised three sources and six detectors, yielding up to 18 within-module channels together with a variable number of between-module channels, with source–detector separations spanning 8–60 mm. For the present study, one module over each hemisphere was used to provide bilateral coverage of the dlPFC (left and right dlPFC).

#### 2.5.2 TD-fNIRS preprocessing

Shared Near Infrared Spectroscopy Format (SNIRF) files were exported from the Kernel Flow 2 system using the manufacturer’s Hb Moments pipeline (Dubois et al., 2024, see software section), which yields oxygenated (HbO) and deoxygenated (HbR) haemoglobin concentration changes in channel space. Each recording was trimmed to the relevant task period, optode channels removed that had with insufficient or atypically shaped photon time-of-flight distributions, computes the moments of the time-of-flight distribution, converts these to HbO and HbR concentration changes, and applies motion-artefact correction and global signal regression (the latter by regressing out the mean signal across short-separation intra-module channels). Full pipeline details are provided by the manufacturer (Dubois et al., 2024). On the resulting HbO time series, we applied a band-pass filter (0.02–0.2 Hz) to isolate the task-relevant haemodynamic frequency band, and restricted subsequent task-evoked analyses to long-separation channels overlying the left and right dlPFC (l-dlPFC, r-dlPFC respectively).

#### 2.5.3 dlPFC task-evoked haemodynamic analysis

Task-related haemodynamic responses were estimated using a general linear model (GLM), implemented with MNE-NIRS (Luke et al., 2021) and Nilearn (Abraham et al., 2014), fitted to dlPFC HbO time series across the full task-battery recording. All cognitive tasks in the battery (Immediate Object Recall, Digit Span, 2N-Back, Stroop, and Delayed Object Recall) were modelled as regressors of interest using a finite impulse response (FIR) basis spanning a 20-second post-stimulus window, allowing the task-evoked response to be estimated without assuming a fixed response shape. Practice blocks and the immediate object-memory encoding period were included as nuisance regressors, and the leading 10 principal components of the short-separation channels (per chromophore) were added to account for systemic and superficial haemodynamic contributions. Although the manufacturer pipeline already regresses out the mean short-separation signal, this captures only the average superficial component; the principal-component regressors were included to account for additional structured physiological variance not removed by mean regression. Low-frequency drift was modelled with a discrete-cosine drift basis. For each session, peak task-evoked activation (β max) was subsequently extracted for the 2N-Back and Switching Stroop conditions only.

#### 2.5.4 Channel coverage and coverage scaling

Channel coverage for each session was defined as the percentage of channels within the prefrontal module that retained usable signal after channel-level quality checks. Because estimated response amplitude fell as coverage increased, amplitudes were rescaled as a function of session coverage. Scaling was applied after the GLM described in 2.5.3. The fitted HbO FIR coefficients of each session, for every channel and delay, were multiplied by a session-specific scale factor s(c), where c is the session’s prefrontal coverage (%), before peak and maximum gradient amplitudes were extracted. For sessions with coverage below a threshold c_thresh, the scale factor was

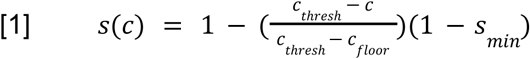

and sessions at or above c_thresh were left unchanged (s = 1). The factor falls linearly as coverage decreases and reaches a minimum of s_min at c_floor, a reference coverage set one percentage point below the lowest coverage observed in the fitting data. As the GLM is linear in the signal, this is equivalent to scaling the HbO signal before model fitting.

The minimum scale s_min was defined as the mean amplitude of sessions with c ≥ c_thresh divided by the mean amplitude of the lowest-coverage 10% of sessions. Amplitude here was the bilateral dlPFC peak response (left plus right), pooled across tasks and taken from the unscaled data.

Parameters c_floor and s_min were estimated using five-fold cross-validation, with folds split by participant. In each fold they were derived from the training participants only, and the mean across folds was used for the final correction applied to all sessions. This yields parameter estimates that are not dominated by any single subset of participants. The threshold c_thresh was selected by evaluating candidate values from 50% to 100% and balancing the reduction in the correlation between amplitude and coverage against the preservation of test-retest reliability (ICC). The final parameters were c_thresh = 90%, c_floor = 28.0% and s_min = 0.305

#### 2.5.5 dlPFC response and cognitive performance association

Associations between task-evoked dlPFC activation and cognitive performance were assessed at the participant level. Neural and behavioural measures were averaged across available scanning sessions for each participant, yielding one observation per participant. Pearson correlations (r) were then computed between log-transformed activation and each behavioural metric, separately for each region of interest (l-dlPFC; r-dlPFC) and task (2N-Back; Switching Stroop). Regression lines were fitted using ordinary least squares. Statistical significance was assessed using two-tailed tests, with thresholds of p < 0.05, p < 0.01 and p < 0.001.

#### 2.5.6 Hemispheric laterality index

To quantify the relative engagement of l- and r-dlPFC, a laterality index (LI) was computed for each task and scan using the standard formulation (Seghier, 2008):

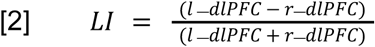

where l- and r-dlPFC are the peak activations (β max) in the left and right hemispheres, respectively. The index is bounded between −1 and +1: +1 denotes exclusively left-hemisphere activation, −1 exclusively right-hemisphere activation, and values near zero balanced bilateral engagement. LI was then examined as a function of age group, sex at birth, and task.

#### 2.5.7 dlPFC response and demographic / lifestyle prediction

Wearable features entered the models in trait-state pairs. Each pair combined a trait feature with a state feature of the same metric: baseline with day deviation, baseline with MACD, slope with recent slope, RMSSD with recent RMSSD, and CV with recent CV (see 2.3.3). Because the models required complete data at every scan, analyses were restricted to participants with complete wearable features and dlPFC targets at all three scans: 36 participants (108 scans; Garmin 17, Whoop 10, Oura 7, Apple 1, Fitbit 1).

Feature selection was performed by exhaustive search over the wearable feature pairs alone. Each candidate model contained one feature pair from each of the three clusters (see 2.3.5), and every possible combination was evaluated with ridge regression (α = 31). Candidates were scored by repeated participant-grouped cross-validation. Each repeat was a 5-fold partition in which all scans from a given participant fell in the same fold, so that no participant appeared in both training and test sets. Features were standardised within each training fold. Within each repeat, out-of-fold predictions from all five folds were concatenated and a single pooled coefficient of determination (R²) was computed against the observed values, rather than averaging fold-wise R². This was repeated across 100 random participant groupings, and the mean pooled R² across repeats was taken as the score.

The search was run separately for bilateral dlPFC peak activation in the 2N-Back and Switching Stroop tasks. Of the two best-scoring models, we retained the one with the best R² across both tasks, so that a single wearable feature set was shared across all targets and both tasks. The selected set comprised the baseline and MACD of REM sleep duration, the CV and recent CV of total calories burned, and the baseline and MACD of respiration rate during sleep.

Demographic variables and confounds were not entered into the search. The wearable feature set was optimised on its own, and these covariates were added afterward to preserve their interpretation as separate contributions. Feature selection was performed once on the full cohort rather than nested within the cross-validation used to evaluate the final models. Because the search was optimised for bilateral dlPFC activation, predictive performance for these targets may be optimistic relative to the other targets, and comparisons between targets should be interpreted with this selection bias in mind.

Ridge regression was then used to predict a range of session-level outcomes (Figure 4b). For each task (2N-Back and Switching Stroop), separate models were fitted for neural targets (l-dlPFC, r-dlPFC and bilateral dlPFC peak activation, the last defined as the sum of left and right peak activation), behavioural targets (summary score and median RT) and composite targets dividing a behavioural measure by bilateral activation (median RT / bilateral activation, RT & dlPFC; summary score / bilateral activation, Perf & dlPFC).

**Figure 3.**
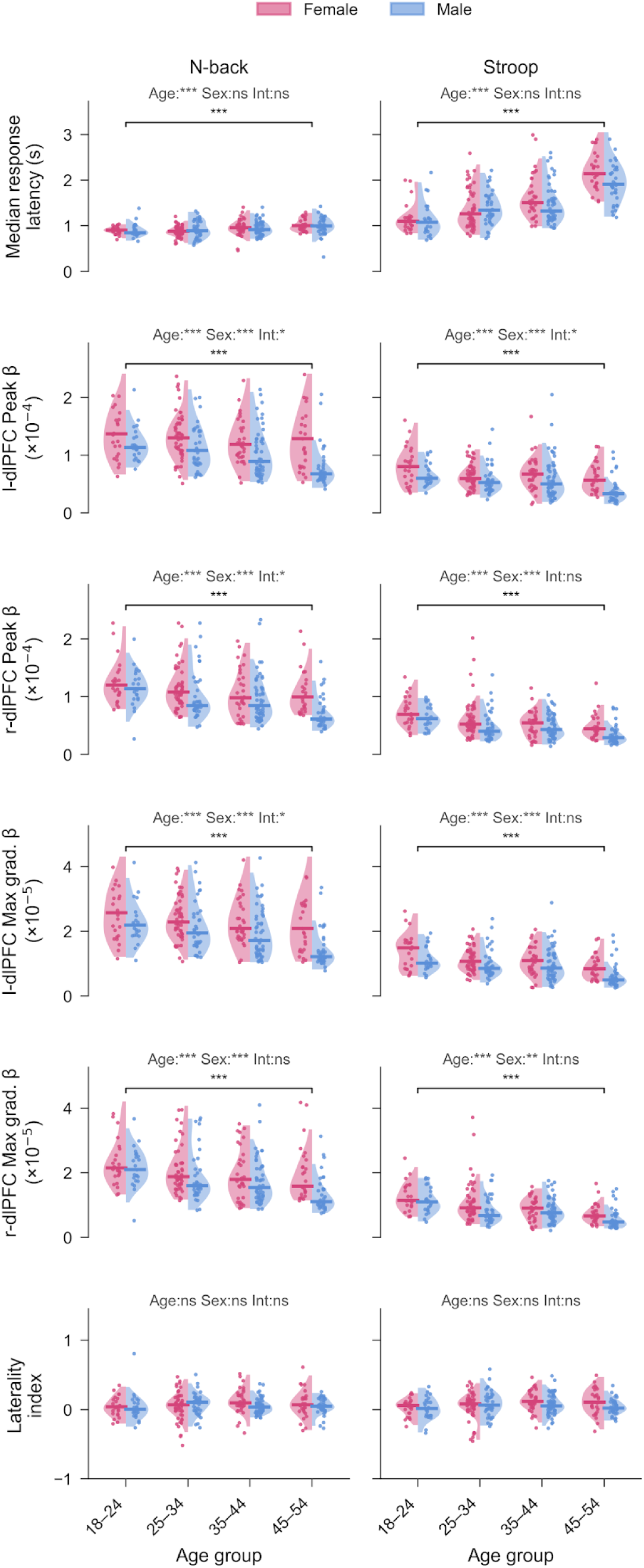
Prefrontal haemodynamic response, response latency and hemispheric laterality by age and sex. Median response latency for the 2N-Back (all responses) and Switching Stroop (all trials) (top row), peak β (×10⁻⁴) for l-dlPFC (second row) and r-dlPFC (third row), maximum gradient β (×10⁻⁵) for l-dlPFC (fourth row) and r-dlPFC (fifth row), and the hemispheric laterality index (LI = (L − R)/(L + R), where L and R are peak β in the left and right hemispheres; bottom row), shown across all scanning sessions (N = 98 participants, 287 scan observations). Each panel shows the distribution split by age group (x-axis: 18–24, 25–34, 35–44, 45–55 years) and sex (Female, pink; Male, blue). Half-violins show the kernel density estimate for each sex within each age group, points are individual scan observations with horizontal jitter, and medians are indicated by horizontal lines within each half-violin. Omnibus age-group differences were tested with Kruskal–Wallis H, and significant effects are marked with a bracket above the relevant panel. Two-way ANOVA (age × sex) statistics are shown above each panel (Age, Sex, Int, for the interaction; *p < 0.05, **p < 0.01, ***p < 0.001; ns, not significant). In contrast to activation magnitude, laterality showed no age-group, sex or interaction effects for either task. Positive LI values indicate left-hemisphere dominance and negative values right-hemisphere dominance.

**Figure 4.**
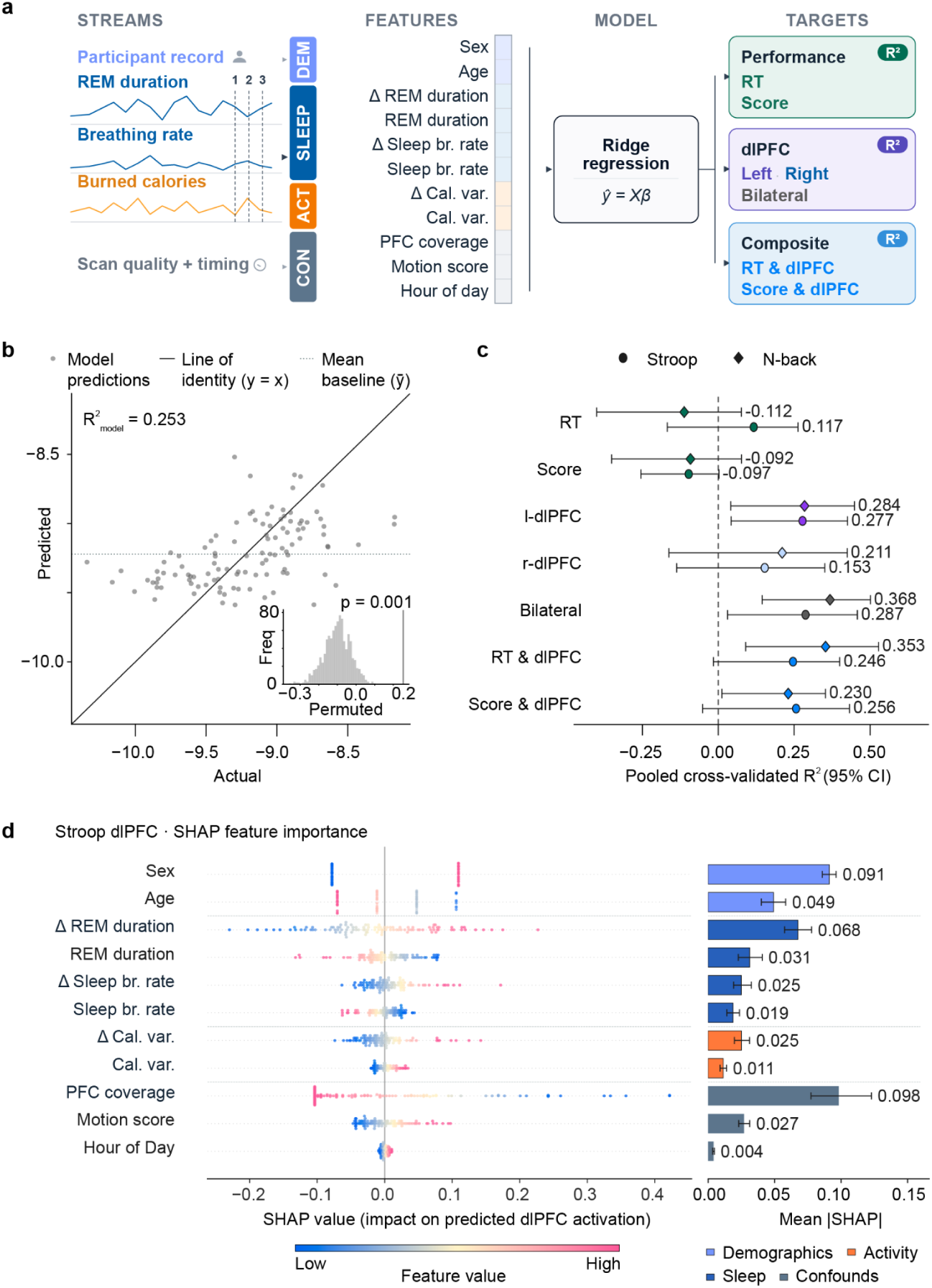
dlPFC activation is better predicted than task performance from demographic, lifestyle and scan-quality features. **(a)** Schematic of the ridge-regression prediction framework. Illustrative wearable traces (left; REM sleep duration, respiration rate during sleep, daily calories burned; ∼120 days, three scans marked) are summarised into features at each scan alongside the participant record and scan-quality data. Ridge regression predicts behavioural (median RT, score), neural (left, right, bilateral dlPFC peak activation) and composite (behavioural ÷ bilateral activation) targets from demographics, sleep and activity features (each a trait–state pair) and scan-quality/temporal confounds. **(b)** Model validation for the representative target, bilateral dlPFC activation during the Switching Stroop (n = 36 participants, 108 scans). Predicted versus actual activation for held-out scans from a single representative cross-validation seed (R² = 0.253) is shown against the line of identity (y = x) and the mean baseline (ȳ), alongside the permutation null distribution from 1,000 shuffles of the target with the observed R² marked (p = 0.001). **(c)** Cross-validated predictive performance for each target in the Switching Stroop (circles) and 2N-Back (diamonds). Markers show the pooled R² from 100 repeats of participant-grouped 5-fold cross-validation in which no participant appears in both the training and test sets and whiskers the 95% confidence interval across repeats; the dashed line marks R² = 0 (no better than the mean baseline). RT, median reaction time; Score, task summary score; l-dlPFC, r-dlPFC and Bilateral, left, right and bilateral dlPFC peak activation; RT & dlPFC, median RT divided by bilateral activation; Score & dlPFC, summary score divided by bilateral activation. **(d)** SHAP feature importance for the bilateral dlPFC Switching Stroop model refitted on the full sample, with features grouped by domain (demographics, sleep, activity, confounds). The beeswarm (left) shows signed SHAP values for every scan, coloured by feature value from low to high; state features are referenced to zero, corresponding to no departure from the participant’s own baseline, and all other features to the cohort mean. The bar plot (right) shows the mean absolute SHAP value per feature, with 95% confidence intervals from 2,000 participant-level bootstrap resamples with the model held fixed.

Neural targets and processing efficiency were log-transformed prior to modelling; behavioural targets and performance efficiency were not. Predictors comprised the six selected wearable features, the demographic covariates age group and sex, and three potential confounding variables: hour of day, head motion, and the number of prefrontal channels retained following signal-quality assessment. Within every cross-validation fold, features were standardised to zero mean and unit variance using parameters estimated from the training data alone, and applied to held-out data to prevent leakage. All models used a fixed ridge penalty (α = 31). Predictive performance was quantified as the mean pooled R² under the same cross-validation scheme as the feature search, and benchmarked against a mean baseline that predicted the training-fold mean for every held-out scan.

Confidence intervals, both for individual-model R² and for between-model differences (ΔR²), were obtained by a participant-level (cluster) bootstrap layered on these 100-repeat predictions. Participants were resampled with replacement 2,000 times, and for each resample the mean pooled R² across the 100 repeats was recomputed on the resampled scans. Model comparisons were evaluated on a common sample of scans complete for both models and matched at the participant level, so that each bootstrap draw yields a paired difference in pooled R². Ninety-five percent confidence intervals were taken as the 2.5th and 97.5th percentiles of the bootstrap distribution, and two-sided bootstrap p-values were defined as in equation 3:

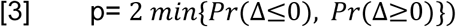

For representative models, above-chance prediction was additionally confirmed by a permutation test. The target was shuffled across scans and a single participant-grouped 5-fold cross-validation was re-run for each of 1,000 permutations. The p-value was defined as (1 + the number of permutations whose pooled R² met or exceeded the observed value) / (1 + 1,000).

#### 2.5.8 Model Interpretability

Feature contributions were quantified with SHAP values (Lundberg and Lee, 2017) from a linear explainer applied to a ridge model refitted on the full dataset, without cross-validation, yielding per-feature attributions for every scan. SHAP values express each feature’s contribution relative to a reference value. For trait features, demographics and confounds, the reference was the cohort mean. For state features, which are expressed as departures from the participant’s own baseline (see 2.3.3), the reference was a value of zero, corresponding to no departure. Their contributions therefore reflect the effect of a participant deviating from their own norm, rather than from the cohort. Mean absolute SHAP values ranked features by overall contribution, and signed SHAP values characterised the direction of each feature’s effect on the predicted target. Features were grouped into demographic, sleep, activity, and confound domains for summary.

### 2.6 Statistical Analysis

Test-retest reliability of dlPFC β estimates and cognitive performance metrics was assessed using a two-way random-effects, absolute-agreement, single-measurement intraclass correlation coefficient (ICC(A,1)), computed with Pingouin (Vallat, 2018). Participants were treated as targets and sessions, ordered by acquisition time, as raters. Because the ICC requires a balanced design, only participants with at least three sessions were included, and participants with more than three were truncated to their first three sessions by acquisition time, yielding three measurements per participant. For each measure we report the ICC(A,1) with its F statistic, p value, and 95% confidence interval, alongside between-person variance (the variance of participant means), within-person variance (the mean of participants’ across-session variances), and their ratio.

Associations between prefrontal activation and cognitive performance were examined using Pearson correlations (r) at the participant level, with neural and behavioural measures averaged across scanning sessions to yield one observation per participant. Correlations between log-transformed activation and behavioural metrics (median RT and accuracy) were computed separately for each region (l-dlPFC; r-dlPFC) and task (2N-Back; Switching Stroop) combination.

Age-related differences in mean peak β and cognitive performance were examined using Kruskal-Wallis H tests across four age groups, with effect size reported as:

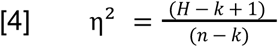

Where a significant omnibus effect was observed, post-hoc pairwise comparisons were conducted using Mann-Whitney U tests with Bonferroni correction. Sex differences were evaluated using Mann-Whitney U tests. Two-way ANOVA (age group × sex) was used to assess main effects and interactions, with partial η² reported as a measure of effect size.

Within-participant changes in β across consecutive scanning sessions were assessed using Wilcoxon signed-rank tests on paired observations, with Bonferroni correction applied across the number of pairwise scan comparisons. All tests were two-tailed. Significance thresholds were set at p < 0.05 (*), p < 0.01 (**), and p < 0.001 (***).

### 2.7 Software

Data were stored and automatically processed using the Connectome Health Platform (https://connectome.health/). Data acquisition and preprocessing were performed using the Kernel Flow 2 data acquisition graphical user interface and the Hb Moments pipeline (Dubois et al., 2024; https://docs.kernel.com/docs/data-export-pipelines). Subsequent analyses were conducted in Python 3.14.2, using NumPy, SciPy, and pandas for data handling, scikit-learn (Pedregosa et al., 2011) for ridge regression, Pingouin (Vallat, 2018) for ICC, and SHAP (Lundberg and Lee, 2017) for feature attribution. GLM modelling was conducted using MNE-NIRS (Luke et al., 2021) and Nilearn (Abraham et al., 2014). Statistical analyses were performed using SciPy and Pingouin.

## 3 Results

### 3.1 Participant characteristics

The LUCID dataset comprised 98 healthy adults (47 female) spanning four age groups: 18–24 years (n = 16, 50% female), 25–34 years (n = 31, 58% female), 35–44 years (n = 30, 40% female), and 45–55 years (n = 21, 43% female). They completed three in-person scanning sessions. Mean inter-session interval was 10.6 ± 5.8 days. Each session yielded simultaneous TD-fNIRS neuroimaging, cognitive task performance (2N-Back and Switching Stroop), pre-scan survey data, and longitudinal wearable-derived lifestyle metrics. Full participant flow and session completion rates are shown in Figure 1.

### 3.2 dlPFC peak activation provides a stable, individual-sensitive functional neural marker

Log-transformed peak dlPFC activation (log β max; Figure 2e) was negatively associated with median response latency for the Switching Stroop in both hemispheres (l-dlPFC r = −0.32; r-dlPFC r = −0.38; both p < 0.01), such that participants with larger prefrontal haemodynamic responses responded faster on average (Figure 2f). The same pattern held for maximum gradient (l-dlPFC r = −0.46; r-dlPFC r = −0.48; both p < 0.001). These associations were not significant for the 2N-Back (peak r = −0.14 and −0.10; maximum gradient r = −0.17 and −0.13), indicating that the coupling between prefrontal recruitment and response speed is clearest under the higher cognitive-control demand of the Switching Stroop.

Importantly, task-evoked dlPFC activation showed evidence of stable individual differences across repeated sessions, alongside within-person variability. Decomposing the variance of all neural and behavioural measures into between- and within-person components (Table 3) showed that between-person variance exceeded within-person variance for every measure (all between:within ratios > 1), with moderate-to-good reliability overall (ICC range 0.54–0.74). Among the neural measures, the bilateral composites were the most reliable (2N-Back bilateral ICC = 0.694, ratio 2.27:1; Stroop bilateral ICC = 0.627, ratio 1.67:1), and aggregating across hemispheres improved stability relative to either side alone (bilateral 2N-Back ICC = 0.694 vs unilateral 0.578–0.609). The predominance of between-person variance suggests that individual differences in peak dlPFC activation are relatively consistent over the short-term assessment period, although day-to-day physiological and contextual factors may also contribute to variation in the measured response.

**Table 3.** ICC and variance components for neural and behavioural measures.

|  | Metric | ICC | F | p | 95% CI | Var<br>(between) | Var (within) | Ratio |
| --- | --- | --- | --- | --- | --- | --- | --- | --- |
| **Stroop** | **Median RT** | 0.740 | 16.27 | $3.01 \times 10^{-54}$ | [0.46, 0.86] | 0.2237 | 0.08278 | 2.70:1 |
| | **l-dIPFC $\beta$ max** | 0.577 | 5.201 | $1.93 \times 10^{-21}$ | [0.47, 0.68] | $4.54 \times 10^{-10}$ | $3.342 \times 10^{-10}$ | 1.36:1 |
| | **r-dIPFC $\beta$ max** | 0.568 | 5.012 | $1.39 \times 10^{-20}$ | [0.45, 0.67] | $3.739 \times 10^{-10}$ | $2.853 \times 10^{-10}$ | 1.31:1 |
| | **dIPFC bilat. $\beta$ max** | 0.627 | 6.205 | $1.01 \times 10^{-25}$ | [0.52, 0.72] | $1.491 \times 10^{-9}$ | $8.935 \times 10^{-10}$ | 1.67:1 |
| | **l-dIPFC max grad. $\beta$** | 0.627 | 6.235 | $7.60 \times 10^{-26}$ | [0.52, 0.72] | $1.395 \times 10^{-11}$ | $8.364 \times 10^{-12}$ | 1.67:1 |
| | **r-dIPFC max grad. $\beta$** | 0.67 | 7.506 | $1.01 \times 10^{-30}$ | [0.57, 0.76] | $1.358 \times 10^{-11}$ | $6.749 \times 10^{-12}$ | 2.01:1 |
| | **dIPFC bilat. max grad. $\beta$** | 0.705 | 8.712 | $7.02 \times 10^{-35}$ | [0.61, 0.78] | $5.06 \times 10^{-11}$ | $2.143 \times 10^{-11}$ | 2.36:1 |
| **2N-Back** | **Median RT** | 0.538 | 5.229 | $2.40 \times 10^{-21}$ | [0.39, 0.66] | 0.01444 | 0.01306 | 1.11:1 |
| | **l-dIPFC $\beta$ max** | 0.578 | 5.070 | $7.52 \times 10^{-21}$ | [0.46, 0.68] | $1.72 \times 10^{-9}$ | $1.255 \times 10^{-9}$ | 1.37:1 |
| | **r-dIPFC $\beta$ max** | 0.609 | 5.648 | $2.14 \times 10^{-23}$ | [0.50, 0.71] | $1.277 \times 10^{-9}$ | $8.195 \times 10^{-10}$ | 1.56:1 |
| | **dIPFC bilat. $\beta$ max** | 0.694 | 7.787 | $9.98 \times 10^{-32}$ | [0.60, 0.77] | $5.486 \times 10^{-9}$ | $2.413 \times 10^{-9}$ | 2.27:1 |
| | **l-dIPFC max grad. $\beta$** | 0.574 | 5.011 | $1.39 \times 10^{-20}$ | [0.46, 0.68] | $5.696 \times 10^{-11}$ | $4.221 \times 10^{-11}$ | 1.35:1 |
| | **r-dIPFC max grad. $\beta$** | 0.634 | 6.147 | $1.73 \times 10^{-25}$ | [0.53, 0.73] | $5.089 \times 10^{-11}$ | $2.933 \times 10^{-11}$ | 1.74:1 |
| | **dIPFC bilat. max grad. $\beta$** | 0.695 | 7.777 | $1.08 \times 10^{-31}$ | [0.60, 0.77] | $1.927 \times 10^{-10}$ | $8.462 \times 10^{-11}$ | 2.28:1 |

Together, the association with Stroop response latency and the cross-session reliability of peak activation and maximum gradient suggest that these measures capture relatively consistent individual differences in prefrontal haemodynamic function. These findings support their potential utility as repeatable, behaviourally relevant functional neural markers.

### 3.3 Variation in peak dlPFC activation with age group and sex at birth

Peak dlPFC activation (β max) was analysed at the scan-observation level (n = 98 participants, 287 observations; Figure 3) using two-way ANOVA on log-transformed β with age group and sex at birth as factors, fitted separately for each of the four region–task combinations. β max differed significantly across age groups in all four region–task combinations (l-dlPFC 2N-Back: F(3,279) = 6.41, p < 0.001, partial η² = 0.065; l-dlPFC Stroop: F(3,279) = 11.25, p < 0.001, partial η² = 0.108; r-dlPFC 2N-Back: F(3,279) = 8.98, p < 0.001, partial η² = 0.088; r-dlPFC Stroop: F(3,279) = 16.04, p < 0.001, partial η² = 0.147). The effect was consistent across hemispheres and tasks, with larger effect sizes for Stroop than for the 2N-Back. In every region–task combination, peak activation was lower in older age groups, decreasing across the age range in both hemispheres. Effect sizes were medium (partial η² = 0.065–0.147), indicating that age group accounted for a meaningful but modest proportion of scan-level variance in activation. Maximum gradient showed the same age and sex effects (all age p < 0.001, all sex p < 0.01).

Within-group variance was substantial across all comparisons, with individual differences dominating over demographic trends. Age group was retained as a covariate in all downstream predictive models. Sex at birth also had a significant main effect in every region–task combination, with higher activation in females (l-dlPFC 2N-Back: F(1,279) = 26.18, p < 0.001, partial η² = 0.086; l-dlPFC Stroop: F(1,279) = 20.46, p < 0.001, partial η² = 0.068; r-dlPFC 2N-Back: F(1,279) = 21.08, p < 0.001, partial η² = 0.070; r-dlPFC Stroop: F(1,279) = 13.55, p < 0.001, partial η² = 0.046). A significant age × sex interaction was present for l-dlPFC activation during the 2N-Back (F(3,279) = 3.57, p = 0.015, partial η² = 0.037) and Stroop (F(3,279) = 2.71, p = 0.045, partial η² = 0.028) and for r-dlPFC activation during the 2N-Back (F(3,279) = 3.09, p = 0.028, partial η² = 0.032), but not for r-dlPFC during the Stroop (F(3,279) = 1.49, p = 0.217), indicating that sex-related differences in activation were not uniform across age groups. On the basis of these effects, sex at birth was retained as a covariate in all downstream predictive models.

### 3.4 dlPFC activation, but not task performance, is predictable from demographic, lifestyle and scan-quality features

We next assessed whether task-evoked dlPFC activation, cognitive performance, or their combination could be predicted from demographic and wearable-derived lifestyle features, and which of these targets was most predictable. The clustering procedure (Section 2.3.5) grouped the wearable metrics into three clusters: Sleep Vitals (heart rate variability, resting heart rate and respiration rate during sleep), Sleep Architecture (time asleep, time in bed, and light and REM sleep durations) and Daytime Activity (maximum heart rate and total calories burned) (Supplementary Material A.4). From these, the feature search (Section 2.5.7) selected one metric per cluster, giving six wearable features from sleep respiration rate, REM sleep duration and total calories burned. Because the models required complete wearable data at every scan, these analyses used a stricter subsample than the activation and reliability analyses above: 36 participants (108 scans).

Bilateral dlPFC activation during the Switching Stroop was predicted above chance (mean pooled R² = 0.287 across 100 repeats), which a permutation test confirmed (p = 0.001, the minimum attainable with 1,000 permutations; Figure 4c). Scan-quality confounds contributed substantially to this model, with prefrontal channel coverage the largest single contributor (mean |SHAP| = 0.098); including them accounts for measurement-related variance in activation that demographic and lifestyle features could not otherwise explain. Among participant-level features, sex contributed most (0.091), with higher predicted activation in females, followed by the recent change in REM sleep duration (0.068) and age group (0.049), with lower predicted activation in older age groups (Figure 4d). The two REM sleep features acted in opposite directions: recent increases in REM sleep relative to a participant’s own baseline were associated with higher predicted activation, whereas a higher habitual REM sleep duration (0.031) was associated with lower predicted activation. The remaining wearable features each contributed between 0.011 and 0.025, head motion 0.027 and hour of day 0.004.

Predictive performance varied by target type (Figure 4c). Bilateral activation was the most predictable target in both tasks (Stroop R² = 0.287; 2N-Back R² = 0.368), with left dlPFC activation similarly predictable (Stroop R² = 0.277; 2N-Back R² = 0.284) and right dlPFC activation less so, its confidence intervals including zero (Stroop R² = 0.153; 2N-Back R² = 0.211). Behaviour, by contrast, was poorly predicted: summary scores were not predictable in either task (Stroop R² = -0.097; 2N-Back R² = -0.092), and median RT was modestly predictable for the Stroop (R² = 0.117) but not the 2N-Back (R² = -0.112). Bilateral activation was more predictable than RT for the 2N-Back (ΔR² = 0.49, 95% CI [0.23, 0.79], p = 0.001), but not significantly so for the Stroop (ΔR² = 0.17 [−0.09, 0.47], p = 0.19). Because the feature search was optimised for bilateral activation, this advantage may be partly inflated by selection (Section 2.5.7).

The composite targets inherited the predictability of activation rather than adding to it. Processing efficiency (RT / bilateral activation) did not differ from bilateral activation in either task (Stroop ΔR² = −0.05 [−0.22, 0.12], p = 0.57; 2N-Back ΔR² = −0.01 [−0.15, 0.14], p = 0.91), and exceeded RT only for the 2N-Back (ΔR² = 0.48 [0.24, 0.77], p < 0.001; Stroop ΔR² = 0.13 [−0.07, 0.32], p = 0.18). Performance efficiency (summary score / bilateral activation) exceeded the summary score in both tasks (Stroop ΔR² = 0.36 [0.08, 0.60], p = 0.014; 2N-Back ΔR² = 0.33 [0.09, 0.63], p = 0.009) but never exceeded bilateral activation, falling below it for the 2N-Back (ΔR² = −0.16 [−0.30, −0.04], p = 0.018; Stroop ΔR² = −0.03 [−0.12, 0.03], p = 0.29). Taken together, the predictable signal resides in the haemodynamic response rather than in task behaviour: the composites are predictable because they carry the activation signal, not because brain and behaviour combine to give independent information.

## 4 Discussion

This study combined longitudinal TD-fNIRS imaging of the dlPFC with wearable-derived lifestyle data in a cohort of 98 healthy adults, asking how much of the individual variation in task-evoked prefrontal activation could be accounted for by stable demographic characteristics and modifiable lifestyle factors. Three results define the picture. Peak dlPFC activation showed moderate test-retest reliability across both tasks, with between-person variance exceeding within-person variance for every neural measure. Greater activation was associated with faster response latency during the Switching Stroop task in both hemispheres, whereas no significant association was observed during the 2N-Back task. Demographic, lifestyle and scan-quality features predicted activation out-of-sample at modest but reliably above-chance levels, with sex, recent change in REM sleep and age the strongest participant-level contributors, and the remaining sleep and activity features, spanning habitual patterns and recent departures from them, adding smaller contributions.

Most informatively, these features predicted activation but not task performance, and measures that combined activation with behaviour (activation scaled by RT or by Perf) were no more predictable than activation on its own. Taken together, these findings point not to task performance per se, nor to a combination of activation and behaviour, but to the magnitude of prefrontal activation itself, which we argue reflects a stable, demand-matched capacity for prefrontal recruitment and is the natural target for this class of measurement.

Peak dlPFC activation differed robustly across age groups in every region–task combination, with the largest effects under the Switching Stroop and with older bands showing smaller task-evoked responses in both hemispheres. Hemispheric laterality, by contrast, showed no age effect, so the gradient reflects a roughly proportional reduction in bilateral activation rather than a shift in left–right balance. One possible interpretation is that this pattern reflects age-related differences in vascular responsiveness, with reduced haemodynamic recruitment contributing to smaller task-evoked responses in older participants. Such changes could affect activation magnitude across both hemispheres without necessarily altering laterality. However, TD-fNIRS measurements of HbO do not independently distinguish vascular responsiveness from neural recruitment or neurovascular coupling. The stronger age-related differences observed during the Switching Stroop may reflect greater sensitivity to individual differences under cognitive demand, although the physiological mechanisms underlying this pattern remain to be established.

This pattern runs counter to the dominant account in the cognitive-ageing literature. In older adults (typically ≥60 years), prefrontal activation during demanding tasks is frequently reported to increase or to become more bilateral, long interpreted as compensatory recruitment that offsets declining neural efficiency (Cabeza, 2002; Gonzalez, 2023), though whether such increases are genuinely compensatory rather than nonspecific remains debated (Morcom and Henson, 2018). Our cohort, however, spans 18–55, a range lying largely below the window in which such compensation is described. Consistent with this, an independent adult-lifespan TD-fNIRS study using the same device locates the transition toward compensatory, dedifferentiated recruitment at around the mid-50s (Chekin et al., 2026), at the very upper edge of our range. The direction of our effect is the opposite: activation falls rather than rises across age bands. The same direction held in the predictive model, where older age groups had lower predicted activation after accounting for sex, lifestyle and scan quality (Section 3.4). We therefore favour a vascular account for our results over compensatory reorganisation. Across adulthood, cerebrovascular reactivity falls and neurovascular coupling becomes disrupted (Zimmerman et al., 2021). Both lower the ceiling on the task-evoked haemodynamic response, well before cognitive decline is detectable. On this view, the smaller responses in older bands reflect reduced vascular capacity rather than a failure of compensation. Two caveats apply. First, haemodynamic ageing signals are hard to interpret on their own, because the measured response depends on vascular as well as neural factors. Second, age was a between-participant grouping, so these effects are cross-sectional and cannot separate vascular ageing from cohort differences.

Sex at birth was a robust contributor to prefrontal recruitment. It showed a significant main effect on activation in every region–task combination, with higher activation in females, and entered into significant age × sex interactions in three of the four combinations (Section 3.3). This positions sex as a primary demographic axis of variation in dlPFC activation rather than a nuisance covariate, and the predictive models bore this out, with sex the strongest participant-level contributor to out-of-sample activation (Section 3.4). The effect sits within a mixed literature on sex differences in prefrontal activation (Hill, Laird and Robinson, 2014; Voyer et al., 2021), and its mechanism, whether haemodynamic, anatomical, or task-strategic, cannot be resolved here.

The contribution of wearable-derived sleep and activity features aligns with evidence that habitual sleep and physical activity each shape prefrontal haemodynamics and executive function (Csipo et al., 2021; Naito et al., 2024; Shen et al., 2024). Our design refines this picture by separating each wearable metric into a trait feature, describing the participant’s habitual pattern, and a state feature, describing their recent departure from it. This follows the disaggregation of between-person and within-person effects recommended for repeated-measures data, where a single undifferentiated predictor conflates the two (Curran and Bauer, 2011). The decomposition proved informative. The largest wearable contribution came from a state feature, the recent change in REM sleep duration relative to the participant’s own baseline (mean |SHAP| = 0.068), which exceeded its trait counterpart, habitual REM sleep duration (0.031). The two acted in opposite directions: recent increases in REM sleep were associated with higher predicted activation, whereas a higher habitual REM sleep duration was associated with lower predicted activation. A single undifferentiated REM sleep feature would have blended these opposing associations. This suggests that prefrontal recruitment tracks departures from a person’s own norm as well as their habitual pattern. Read this way, the wearable feature set offers a tractable, if partial, proxy for the exposome spanning both habitual and proximal windows (Sakowski et al., 2024). The contribution should not be overstated, however. Each wearable feature contributed less than sex and less than prefrontal channel coverage, the largest single contributor. Because scan quality must be accounted for before lifestyle effects can be estimated, the wearable contributions reported here are conditional on the confounds rather than standalone predictive gains, and the features were selected on the same cohort used to evaluate them.

These observations are consistent with a capacity account of prefrontal recruitment rather than the classical neural-efficiency account. The classical formulation holds that more capable individuals achieve equal performance with less activation, implying an inverse coupling between activation and performance (Causse et al., 2017). Our data show the opposite pattern during the Switching Stroop task: greater peak activation was associated with faster responding. One possible explanation is that the task-evoked haemodynamic response reflects an individual’s capacity to recruit prefrontal resources under cognitive demand, with larger responses and faster decisions reflecting a common, relatively stable underlying capacity. However, HbO response magnitude reflects the interaction of neural recruitment and vascular physiology, and the present findings cannot distinguish their respective contributions. With this interpretation in mind, the predictive results further suggest that demographic, lifestyle and scan-quality features capture variation in the haemodynamic response more readily than in task performance, while composites combining activation with behaviour were no more predictable than activation alone.

Practically, combining TD-fNIRS with passively collected wearable data may offer a scalable route to functional readouts of prefrontal engagement that do not require laboratory neuroimaging at every timepoint. Framed cautiously, bilateral dlPFC activation is a candidate endpoint rather than a validated marker: its test-retest reliability and modest but above-chance predictability suggest it could serve as a comparator against which clinical or at-risk cohorts are contrasted - for example, populations with documented prefrontal hypoactivation such as ADHD (Peet et al., 2022; Zhang et al., 2023) - or as a repeatable within-individual readout for wellness and cognitive-performance applications, where the comparison of interest is change over time rather than diagnosis. Given that a substantial share of neurological and neuropsychiatric burden is shaped by modifiable exposures accumulating long before clinical thresholds are reached (Steinmetz et al., 2024; Livingston et al., 2024), an upstream, lifestyle-sensitive functional measure is an attractive target. We stress, however, that these are demonstrations of measurability and association in a healthy cohort, not of clinical utility or causal efficacy; any endpoint use would require prospective validation against established functional and clinical measures.

Several limitations qualify these conclusions. (1) Missing data across the wearable streams was substantial, and the predictive models were consequently fitted on a reduced subsample (36 participants, 108 scans, with complete neural and wearable data) rather than the full cohort; the signal nonetheless survived strict participant-level cross-validation, and missingness is tractable through improved compliance protocols and device provisioning in future iterations. (2) Wearable metrics were not derived identically across devices, and inter-device differences in how sleep and activity are computed may have introduced measurement heterogeneity. These differences could introduce additional noise or systematic bias into the predictive models, potentially affecting estimated lifestyle associations. Future studies would benefit from standardised wearable devices or explicit modelling of device-specific effects. (3) fNIRS signal quality depends on hair type and scalp-optode coupling, which vary systematically across individuals and raise both a noise concern and an equity concern, since poorer coupling for some hair types could bias signal recovery. We accounted for this by scaling activation for prefrontal channel coverage and by including coverage and head motion as covariates in the predictive models (Section 2.5.4; Supplementary Material B). However, the influence of scan quality can be reduced rather than fully removed, since stricter correction would also remove legitimate signal, such as age-related vascular differences. (4) The two tasks differ structurally in a way that complicates direct comparison: the 2N-Back is externally paced and advances regardless of response, so its response latencies index decision speed under an imposed deadline, whereas the Switching Stroop is self-paced, so its response latencies index the duration of deliberate conflict resolution. Longer Stroop responses may therefore reflect more thorough resolution rather than slowing, and cross-task comparison of reaction-time-derived indices should be made with this asymmetry in mind. (5) Finally, the feature search was performed on the full cohort and optimised for bilateral activation, so predictive performance for these targets may be optimistic relative to the others; the paired target comparisons were not corrected for multiple testing; the comparison of predictive performance across targets mixes log-transformed and untransformed outcomes (limiting strict R² comparability); and the design is observational and cross-sectional with respect to age. Relatedly, we did not record years of education or other proxies of cognitive reserve, which frequently predict cognitive performance more strongly than chronological age and are known to modulate prefrontal recruitment during executive tasks (Springer et al., 2005; Stern, 2009; Ducharme-Laliberté et al., 2022); unmeasured reserve may therefore partly confound the age- and performance-related effects reported here, and should be captured and modelled in future waves. The consistency of out-of-sample prediction and the convergence on a coherent capacity account strengthen, but cannot substitute for, the causal and longitudinal tests these hypotheses ultimately require.

In sum, individual differences in prefrontal recruitment during demanding cognition are partly structured by who we are and how we live. Sex, recent change in REM sleep and age were the strongest participant-level contributors to predicted dlPFC activation, the remaining sleep and daily-activity features added smaller contributions, and the quantity most legible to these factors was the haemodynamic response itself, reflecting the stable capacity for demand-matched prefrontal recruitment, rather than task performance. We have argued that this capacity plausibly reflects the lifestyle-sensitive, vascularly expressed headroom of the prefrontal haemodynamic response, and that combined fNIRS–wearable measurement offers a practical way to track it. Whether that capacity can be shifted, and whether shifting it matters for cognition and health, are the questions this work sets up rather than settles.

## 5 Funder Information Declared

This work was supported by UK Research and Innovation (UKRI) through UKRI Impact Acceleration Awards EPSRC EP/X52556X/1 and MRC MR/X502959/1, and by Connectome GmbH. Author relationships with the study sponsor are declared under Competing Interests.

## 6 Declarations

### Competing interests

This study was sponsored by Connectome Health (Connectome GmbH), and several authors hold financial and professional interests in the company. [RMH], co-first author and co-investigator, is Chief Science Officer and a co-founder of Connectome Health; [SRS], the principal investigator, is an investor in and advisor to Connectome Health; [DT] is a machine learning engineer at Connectome Health; and [LS] is Chief Executive Officer and a co-founder of Connectome Health. These relationships constitute financial and professional interests in the subject matter of this work. The remaining authors declare no competing interests.

## 7 Author Contributions

**Rufus Mitchell Heggs:** Conceptualisation; study design; ethics submission; participant recruitment, management and scanning; data analysis (Figures 1–4 and Supplementary figures); writing - original draft (introduction, results, discussion). **Daniel Tamkin:** Data analysis: GLM design matrix and parameter selection (Figure 2); wearable feature extraction, feature search and ridge regression analysis (Figure 4); writing: original draft (methods, figure 4 results, discussion). **Tauheed Ibrahim:** Data analysis: identification and development of the prefrontal coverage-scaling correction; update of Figures 2 and 3; provision of RT-fixed and PFC-scaled dlPFC data (Figure 4); writing: original draft (methods: channel coverage and coverage scaling). **Lucas Scherdel:** Conceptualisation; study design; writing: review and editing (introduction, discussion). Data analysis: Feature selection. **Anita Snowdon-Farrell:** Participant recruitment, management and scanning, writing: original draft (methods: study design, setup and participant management). **Alfred Curry:** Data analysis: wearable metric criteria (Supplementary Figures); writing: original draft (methods: selection of wearable metrics) and reviewed text. **Onayomi Rosenior-Patten:** Data acquisition (scanning). **Simon R Schultz:** Conceptualisation; principal investigator; supervision; writing: review and editing (introduction, discussion).

## Notes

### Summary of Updates

The manuscript has been substantially revised to incorporate an expanded cohort of 98 participants, compared with 92 in the original version, and updated statistical analyses. Figure 2 and the associated Results have been revised to clarify the relationship between prefrontal activation and response latency, distinguishing significant associations during the Switching Stroop task from nonsignificant associations during the 2N-Back task. Peak activation and maximum response gradient are now reported separately. Test-retest reliability estimates have been updated, with additional variance analyses presented in Table 3. Figure 3 provides expanded analyses of age and sex differences in prefrontal activation and hemispheric laterality. Figure 4 and the associated Results have been updated to compare the predictability of neural activation, cognitive performance, and combined measures using demographic, wearable-derived lifestyle, and scan-quality features. The Methods have been expanded to clarify haemodynamic preprocessing, signal-quality correction, wearable feature extraction, and predictive modelling. The Abstract and Discussion have been revised to reflect the updated findings and their interpretation. The author list has also been updated to include Tauheed Ibrahim.

