## Supplementary for "Prefrontal activation predicts response latency and is shaped by age and lifestyle"

### Supplementary Material A: Analysis of wearable data

#### A.1: Metric coverage

Not all participants provided us with wearable data successfully and that data was not complete. Of the 98 participants with 3 scans, only 63 had any wearable data reported by Terra. The reporting of metrics was also inconsistent across the metrics, as shown in Figure S1, which shows the coverage of reported wearable metrics before the first scan. From the full list of metrics shown, we selected those that had at least 40 participants with either 10 days reported of the last 30 or 30 reported of the last 90 - see Figure S1. This left us with 19 metrics.

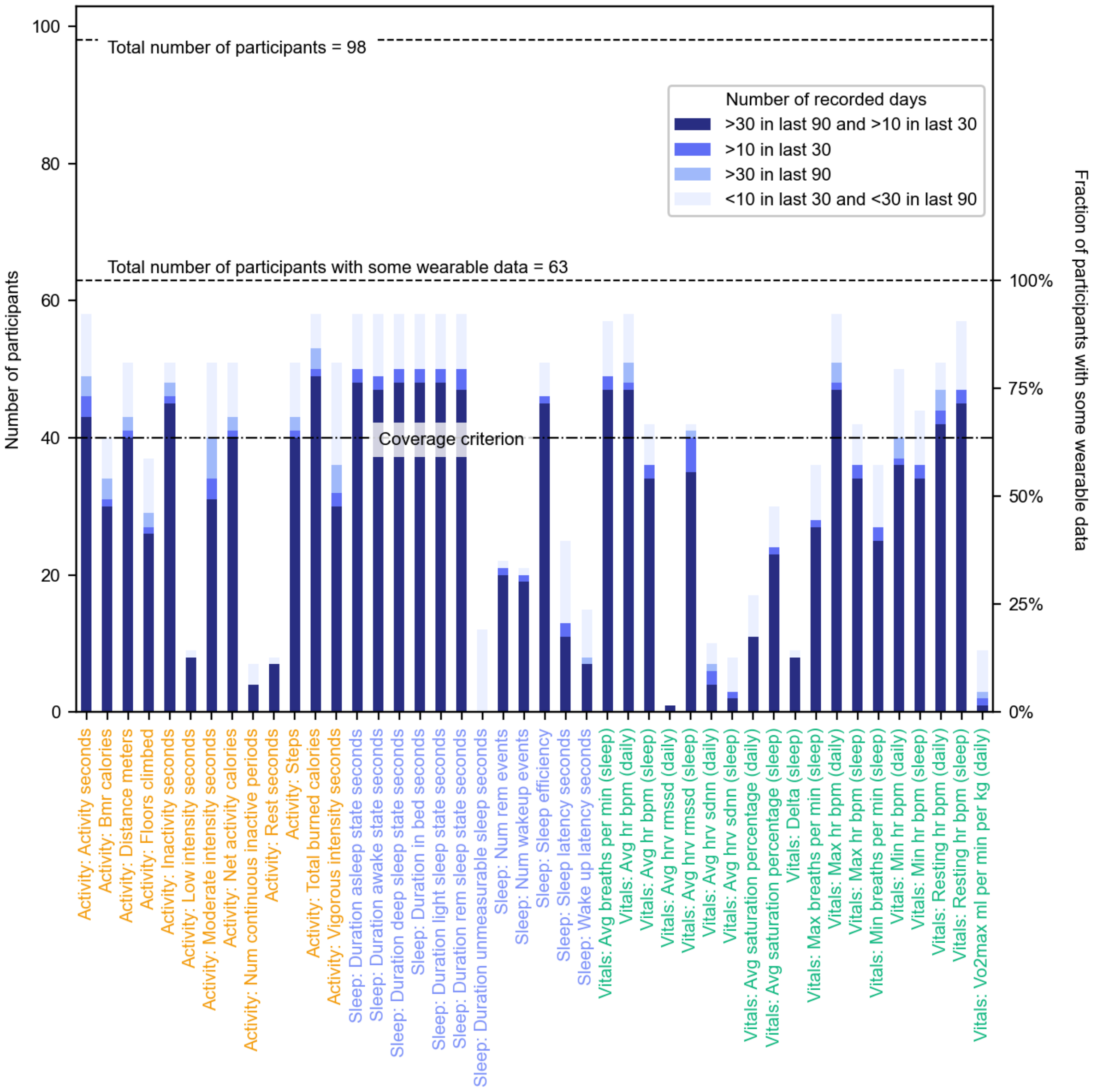


**Figure S1. Wearable metric coverage.** Each bar shows the number of participants in each coverage band for each wearable metric. Coverage bands represent the number of recorded days in the last 90 or 30 days before the participant's first scan.

##### A.2: Coverage across providers

The wearable data came from 6 providers, Apple, Fitbit, Garmin, Google, Oura and Whoop. We wanted metrics that had consistent coverage across providers so limited the features to those that were recorded by the top 4 providers in our sample: Apple, Garmin, Oura and Whoop (Fitbit and Google only had one participant each). This criterion limited the list to 12 metrics: 'Activity: Activity seconds', 'Activity: Total burned calories', 'Sleep: Duration asleep state seconds', 'Sleep: Duration awake state seconds', 'Sleep: Duration deep sleep state seconds', 'Sleep: Duration in bed seconds', 'Sleep: Duration light sleep state seconds', 'Sleep: Duration rem sleep state seconds', 'Vitals: Avg breaths per min (sleep)', 'Vitals: Avg hr bpm (daily)', 'Vitals: Max hr bpm (daily)', 'Vitals: Resting hr bpm (sleep)'. We further excluded 'Activity: Activity seconds' because of known inconsistency of reporting: Whoop reported duration of an activity logged in the app by the user, whereas other providers reported total moving duration in a day. We also included sleeping heart rate variability because it only failed the second criterion due to the fact that Apple report an SDNN measure whereas all other providers use RMSSD. We decided to treat these as equivalent, which was justified in later analysis. This resulted in a list of 12 metrics.

The distribution of the baseline and coefficient of variation (see Section 2.3.3) of these 12 metrics is shown in Figure S2. Many of the features show systematic variation by provider. We quantified this using the Kruskall-Wallis H statistic, the values of which are shown above each panel. We used this as a criterion to exclude the metrics with the worst inter-provider variation. We chose to remove the metric if both the baseline and coefficient of variation showed inter-provider variability with a p-value of <0.001. This was chosen as it was harsh enough to remove the worst metrics, but still kept a reasonable number of them, as few had no inter-provider variability. This means that provider differences remains a caveat to our conclusions. The cut-off resulted in the exclusion of 3 further metrics: 'Sleep: Duration awake state seconds', 'Sleep: Duration deep sleep state seconds' and 'Vitals: Avg HR bpm (daily)'. No significance was detected between the distributions of HRV calculated through different methods, justifying our decision to treat them as equivalent.

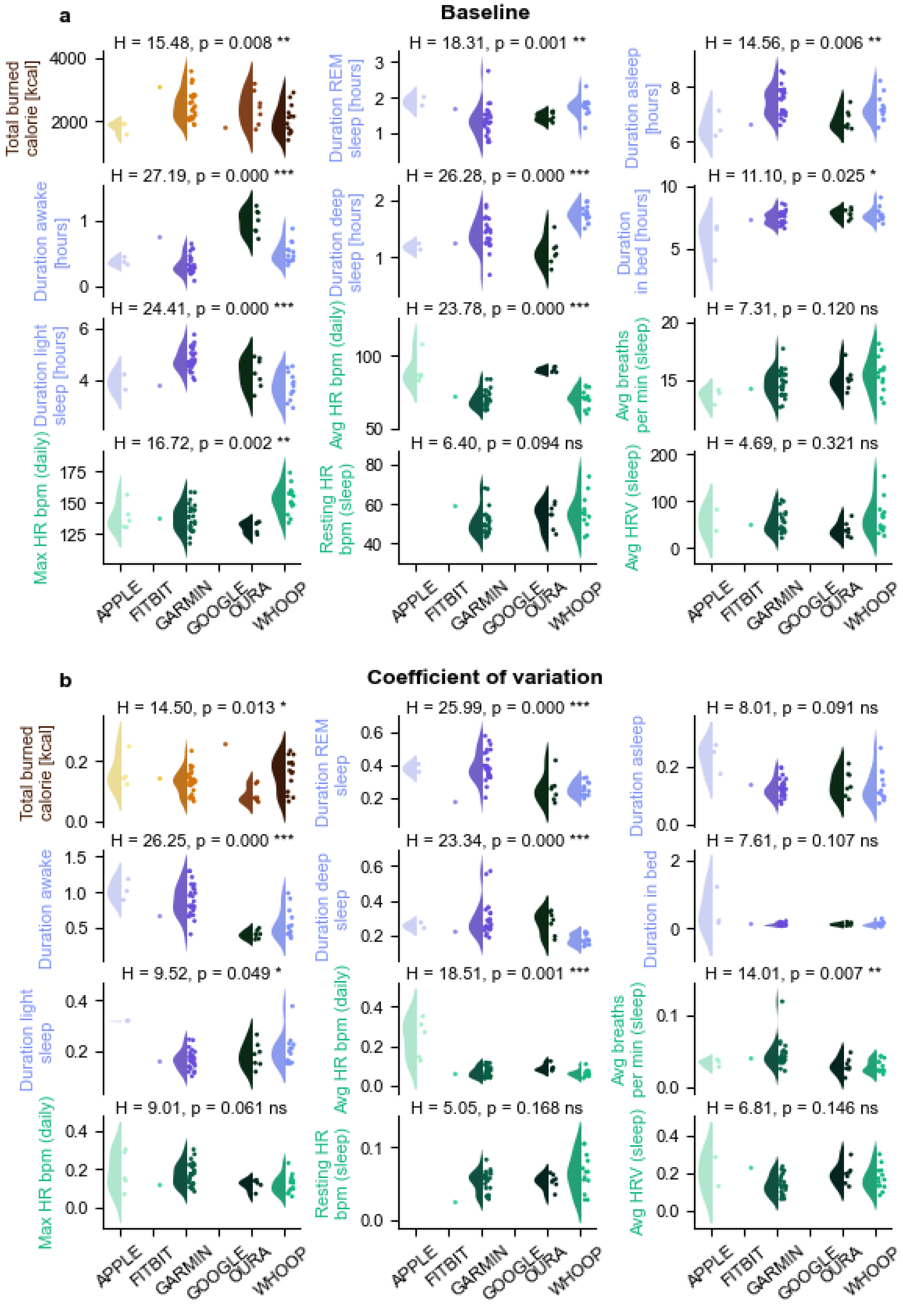


**Figure S2: Wearable metric baselines vary by provider.** Distributions of the baseline value of wearable metrics calculated for the final scan of participants (see Methods), split by the wearable provider. For each plot we show levels of significance of inter-provider difference calculated using a Kruskal-Wallis H-test, along with asterixes showing level of significance .

##### A.3: Within and across participant distributions

Distributions of the 12 metrics selected above are shown in Figure S3a. They show reasonable variation both within and across participants, with peak values, such as 7-8 hours sleep, at sensible values. The activity metric shows an asymmetric, approximately Poisson-shaped distribution, reflecting that on certain days participants might be substantially more active than on most days. Sleep metrics show approximately symmetric, normally distributed values across the sample. One participant's distribution of in bed duration is peaked at 0. This is due to an inconsistency in the recording by Apple Health, where anomalously low values were sometimes recorded (less than the sleep time). This could be rectified by removing anomalous values, however we found that this metric was not a good predictor of dlPFC activity, so did not feel it necessary to correct a single participant.

Lifestyle metrics are only useful measures if they provide different information to demographics. In Figure S3b, we show the distributions of selected metrics broken down by age and sex. While some small trends could be inferred by eye, such as a slight increase sex split in activity, the more significant inference is that the variation within these groups is generally far greater than any difference between groups. This means that the wearable measures can provide us with more information than age or sex alone. Furthermore, it demonstrates that we have successfully sampled a range of activity and sleep levels within these demographic groups. To quantitatively justify this we calculated the ratio between the standard deviation of each group (age or sex) and the difference between the group and total medians, shown Table X ( X Std ratios age sex.xlsx ). For all but the Sleeping HRV age 45-54, the ratio is above 1, inferring that for all other metrics there is useful extra information in the wearable metrics. This makes sense as HRV is known to decline with age [1].

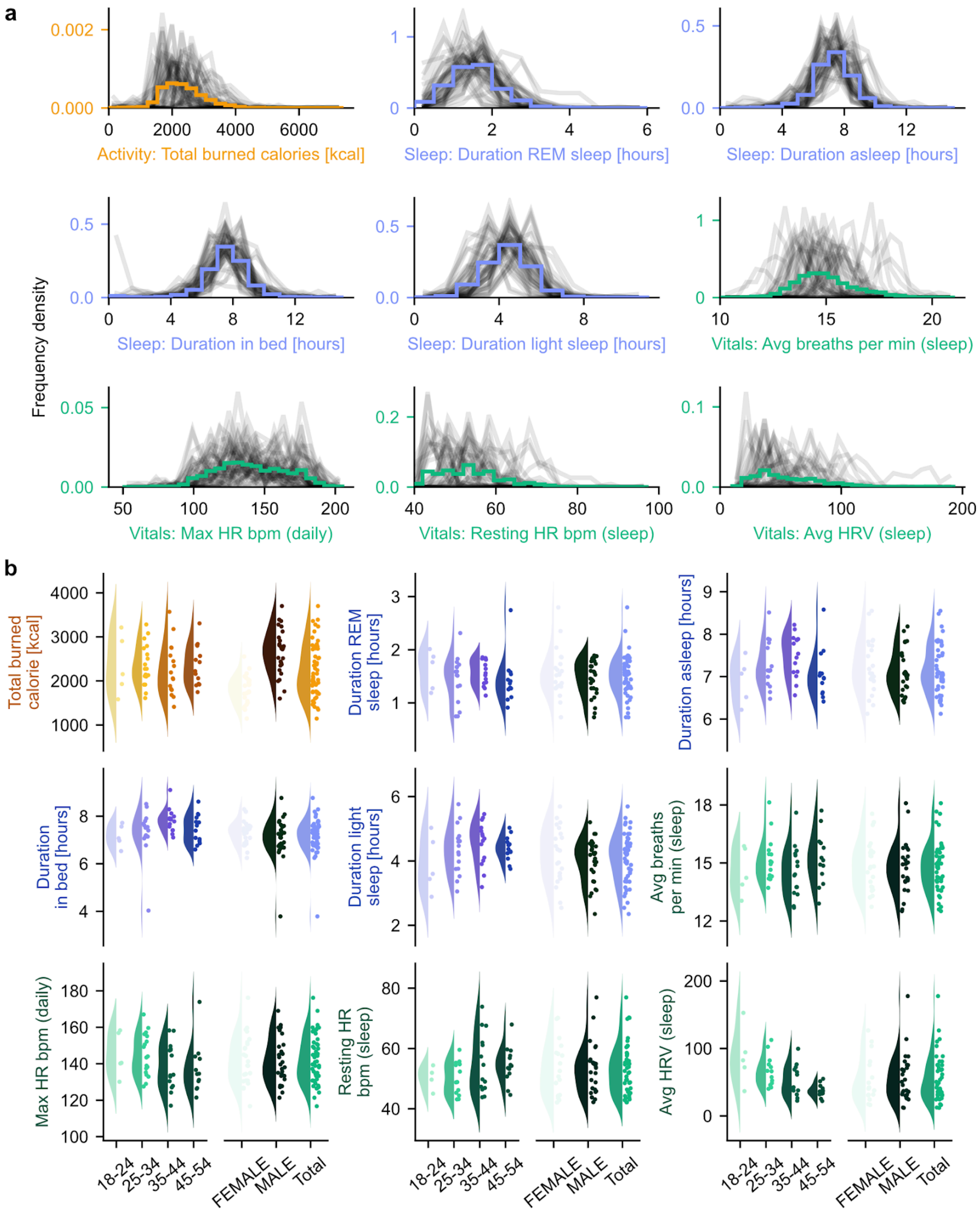


**Figure S3. Wearable metric distributions (a)** Distributions of wearable metrics with adequate coverage (see A.1) across the full sample (coloured bars) and for individual participants (black lines). For most metrics, values cluster around a central peak with meaningful inter-individual variation. **(b)** Distributions of participant's baseline value for the well covered metrics stratified by sex at birth and age group across the full sample. Within-group variance is large relative to any trends across demographic groups, suggesting that wearable-derived digital phenotypes capture individual-level variation beyond what is explained by age or sex alone.

#### A.4 Correlation between metrics

To further select metrics that do not contain redundant information about participants, we investigated the correlation of the metrics with each other. For each participant, we computed the Spearman correlation between each pair of metrics across their daily values, and averaged these across participants after Fisher z-transformation (see Section 2.3.5). A dendrogram showing the result of this clustering algorithm is shown in Figure S4. The dendrogram naturally separates the metrics into 3 clusters that can be intuitively understood. From the bottom of the figure, there is a ‘Daytime Activity’ cluster containing max HR and burned calories, a ‘Sleep Architecture’ cluster containing various sleep duration measures and a ‘Sleep Vitals’ cluster containing heart and respiration stats during sleep. This clustering allowed us to train the ridge regression models using maximally independent wearable features by only allowing one feature from each cluster.

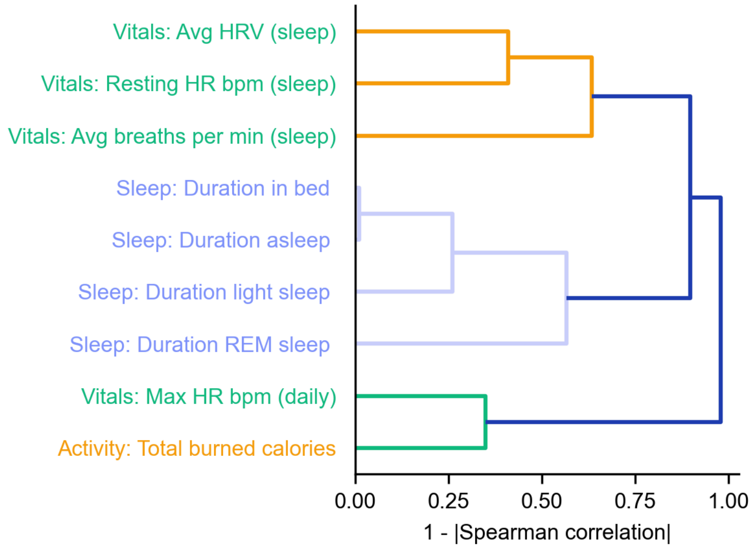


**Figure S4. Wearable data shows correlation within but not across lifestyle domains.** Dendrogram showing the clustering of metrics based on their correlations.

### Supplementary Material B: HbR

#### HbR, Prefrontal coverage scaling

| Metric | ICC | F | p | 95% CI | Var (between) | Var (within) | Ratio |
|---|---|---|---|---|---|---|---|
| **Stroop median RT** | 0.74 | 16.27 | $3.01 \times 10^{-54}$ | [0.46, 0.86] | 0.2237 | 0.08278 | 2.70:1 |
| **Stroop l-dlPFC β max** | 0.603 | 5.681 | $1.54 \times 10^{-23}$ | [0.49, 0.70] | $5.95 \times 10^{-10}$ | $3.933 \times 10^{-10}$ | 1.51:1 |
| **Stroop r-dlPFC β max** | 0.64 | 6.394 | $1.74 \times 10^{-26}$ | [0.54, 0.73] | $6.269 \times 10^{-10}$ | $3.526 \times 10^{-10}$ | 1.78:1 |
| **Stroop dlPFC bilat. β max** | 0.697 | 8.095 | $8.41 \times 10^{-33}$ | [0.60, 0.78] | $2.182 \times 10^{-9}$ | $9.545 \times 10^{-10}$ | 2.29:1 |

| | | | | | | | |
|---|---|---|---|---|---|---|---|
| **Stroop l-dlPFC max grad. β** | 0.604 | 5.796 | $5.00 \times 10^{-24}$ | [0.49, 0.70] | $1.83 \times 10^{-11}$ | $1.215 \times 10^{-11}$ | 1.51:1 |
| **Stroop r-dlPFC max grad. β** | 0.66 | 7.053 | $4.78 \times 10^{-29}$ | [0.56, 0.75] | $2.088 \times 10^{-11}$ | $1.082 \times 10^{-11}$ | 1.93:1 |
| **Stroop dlPFC bilat. max grad. β** | 0.698 | 8.4 | $7.65 \times 10^{-34}$ | [0.60, 0.78] | $6.935 \times 10^{-11}$ | $3.033 \times 10^{-11}$ | 2.29:1 |
| **2N-Back median RT** | 0.538 | 5.229 | $2.40 \times 10^{-21}$ | [0.39, 0.66] | 0.01444 | 0.01306 | 1.11:1 |
| **2N-Back l-dlPFC β max** | 0.613 | 5.71 | $1.16 \times 10^{-23}$ | [0.51, 0.71] | $2.375 \times 10^{-9}$ | $1.494 \times 10^{-9}$ | 1.59:1 |
| **2N-Back r-dlPFC β max** | 0.707 | 8.27 | $2.12 \times 10^{-33}$ | [0.62, 0.78] | $2.302 \times 10^{-9}$ | $9.554 \times 10^{-10}$ | 2.41:1 |
| **2N-Back dlPFC bilat. β max** | 0.729 | 9.051 | $5.63 \times 10^{-36}$ | [0.64, 0.80] | $8.208 \times 10^{-9}$ | $3.043 \times 10^{-9}$ | 2.70:1 |
| **2N-Back l-dlPFC max grad. β** | 0.626 | 5.973 | $9.04 \times 10^{-25}$ | [0.52, 0.72] | $8.391 \times 10^{-11}$ | $5.006 \times 10^{-11}$ | 1.68:1 |
| **2N-Back r-dlPFC max grad. β** | 0.678 | 7.297 | $5.87 \times 10^{-30}$ | [0.58, 0.76] | $7.79 \times 10^{-11}$ | $3.702 \times 10^{-11}$ | 2.10:1 |
| **2N-Back dlPFC bilat. max grad. β** | 0.724 | 8.818 | $3.18 \times 10^{-35}$ | [0.64, 0.80] | $2.855 \times 10^{-10}$ | $1.086 \times 10^{-10}$ | 2.63:1 |

**HbO Supplementary figures**

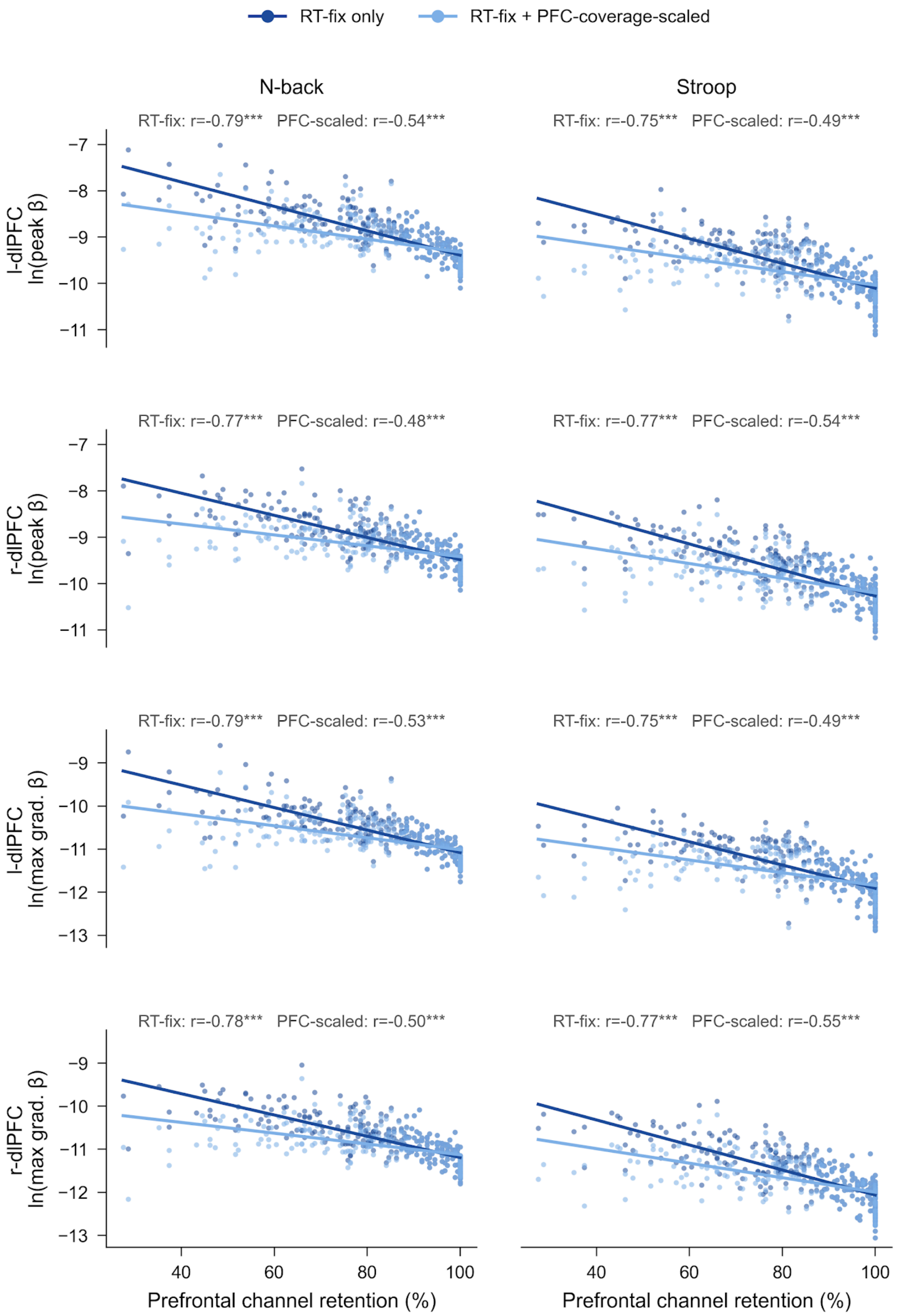


**Supplementary Figure 1.** HbO amplitude against prefrontal coverage for RT fix and PFC scaled data.

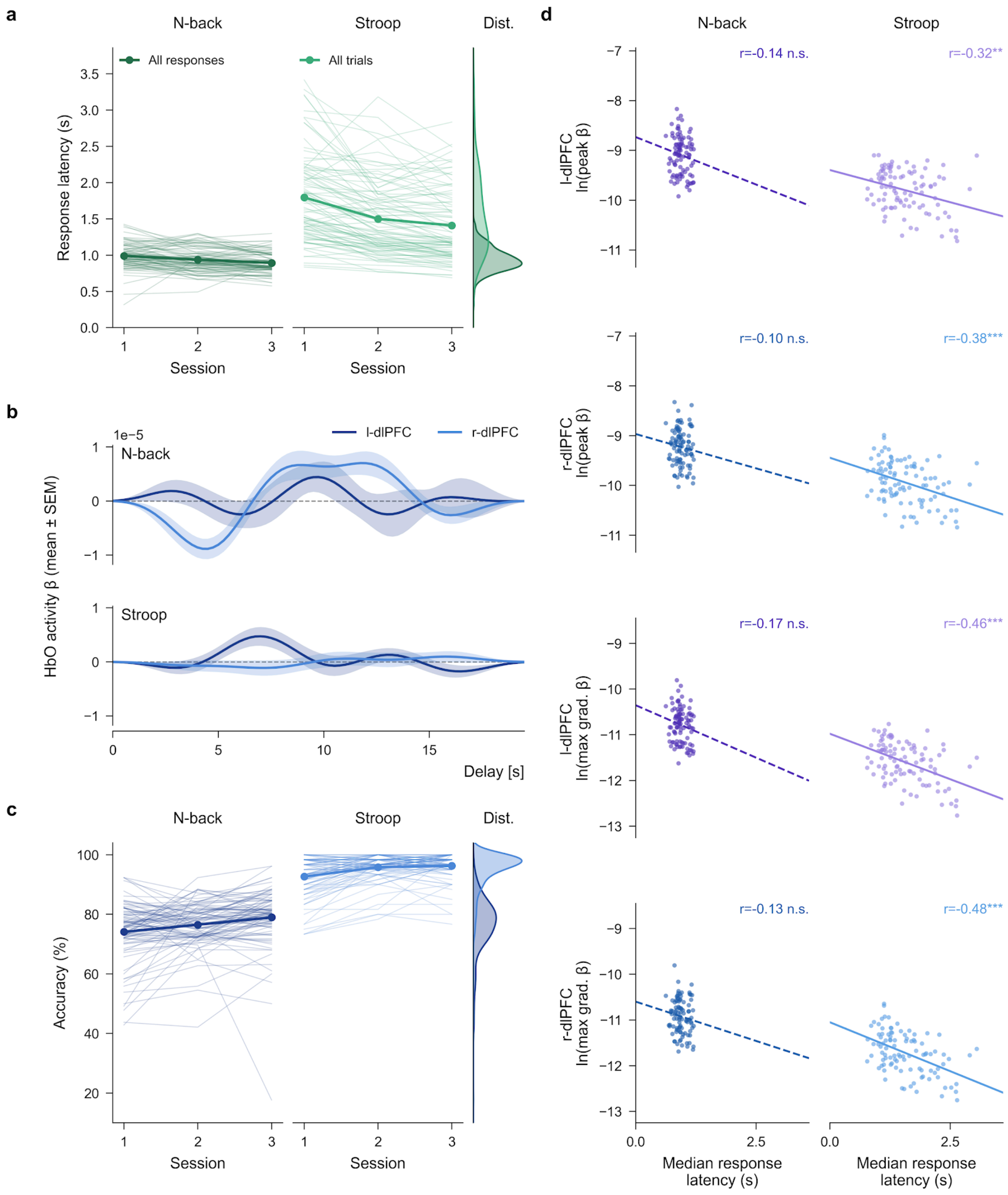


**Supplementary Figure 2.** Response latency, HbO response, task accuracy and latency relationships for PFC scaled data. (a) Response latency across sessions. (b) Mean HbO response. (c) Task accuracy across sessions. (d) Amplitude against response latency.

#### HbR Supplementary figures

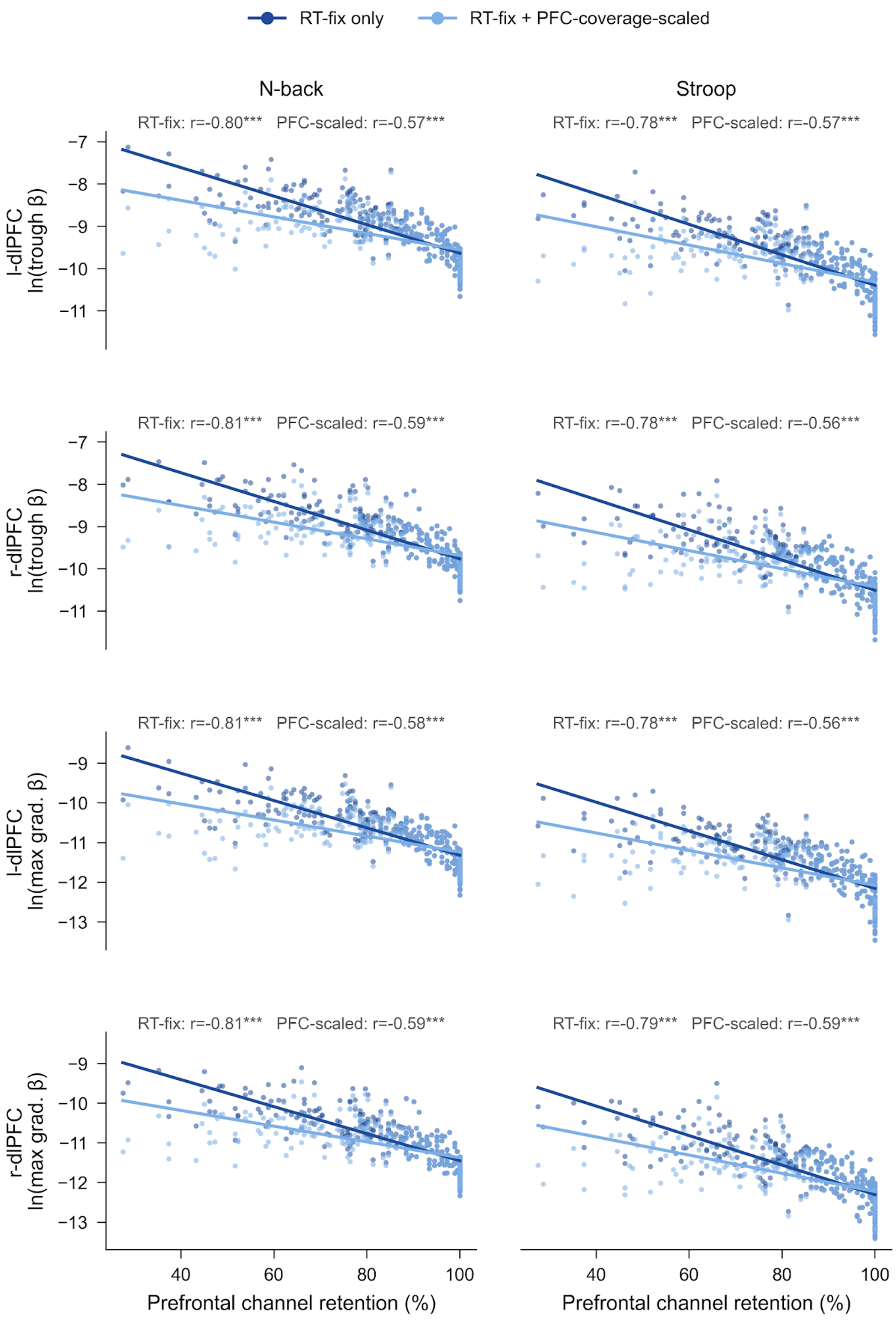


**Supplementary Figure 3.** HbR amplitude against prefrontal coverage for RT fix and PFC scaled data.

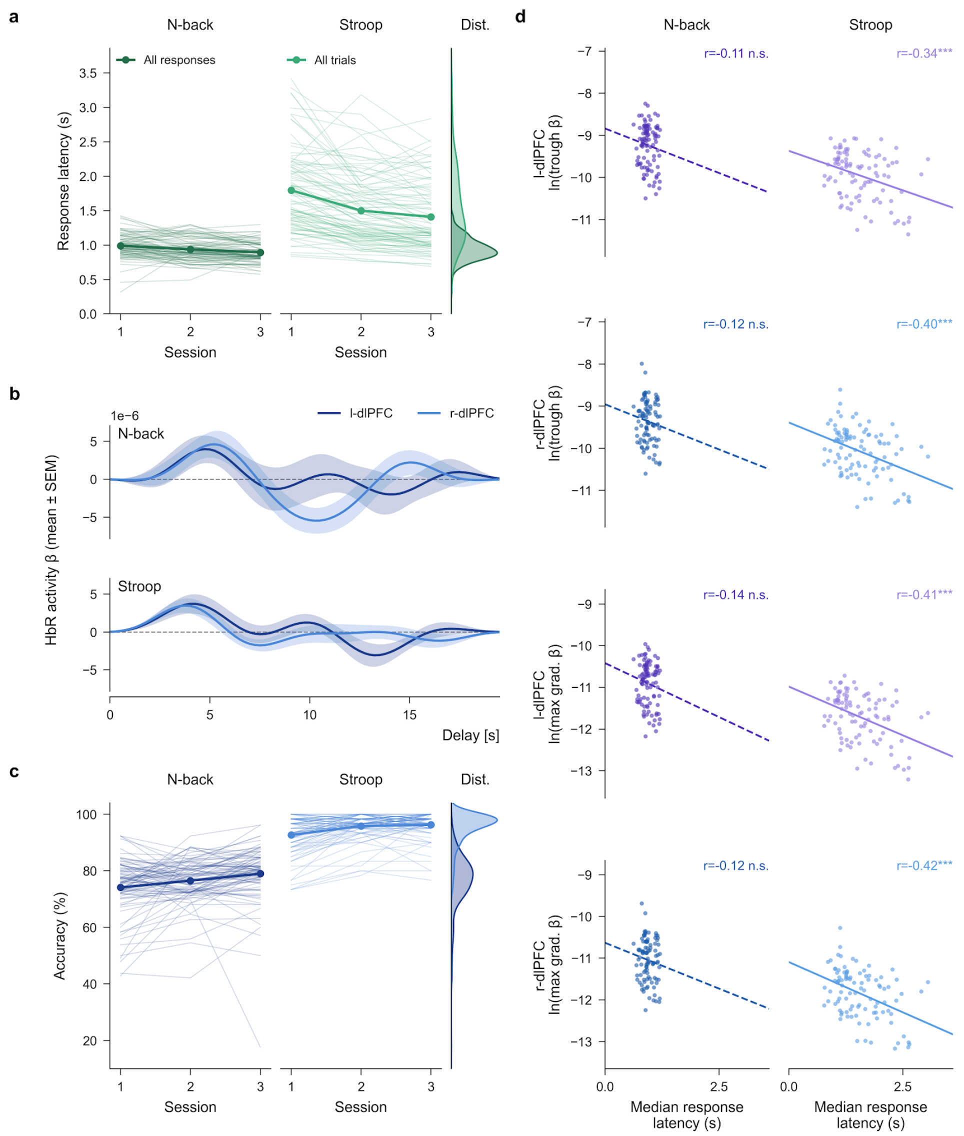


**Supplementary Figure 4.** Response latency, HbR response, task accuracy and latency relationships for PFC scaled data. (a) Response latency across sessions. (b) Mean HbR response. (c) Task accuracy across sessions. (d) Amplitude against response latency.

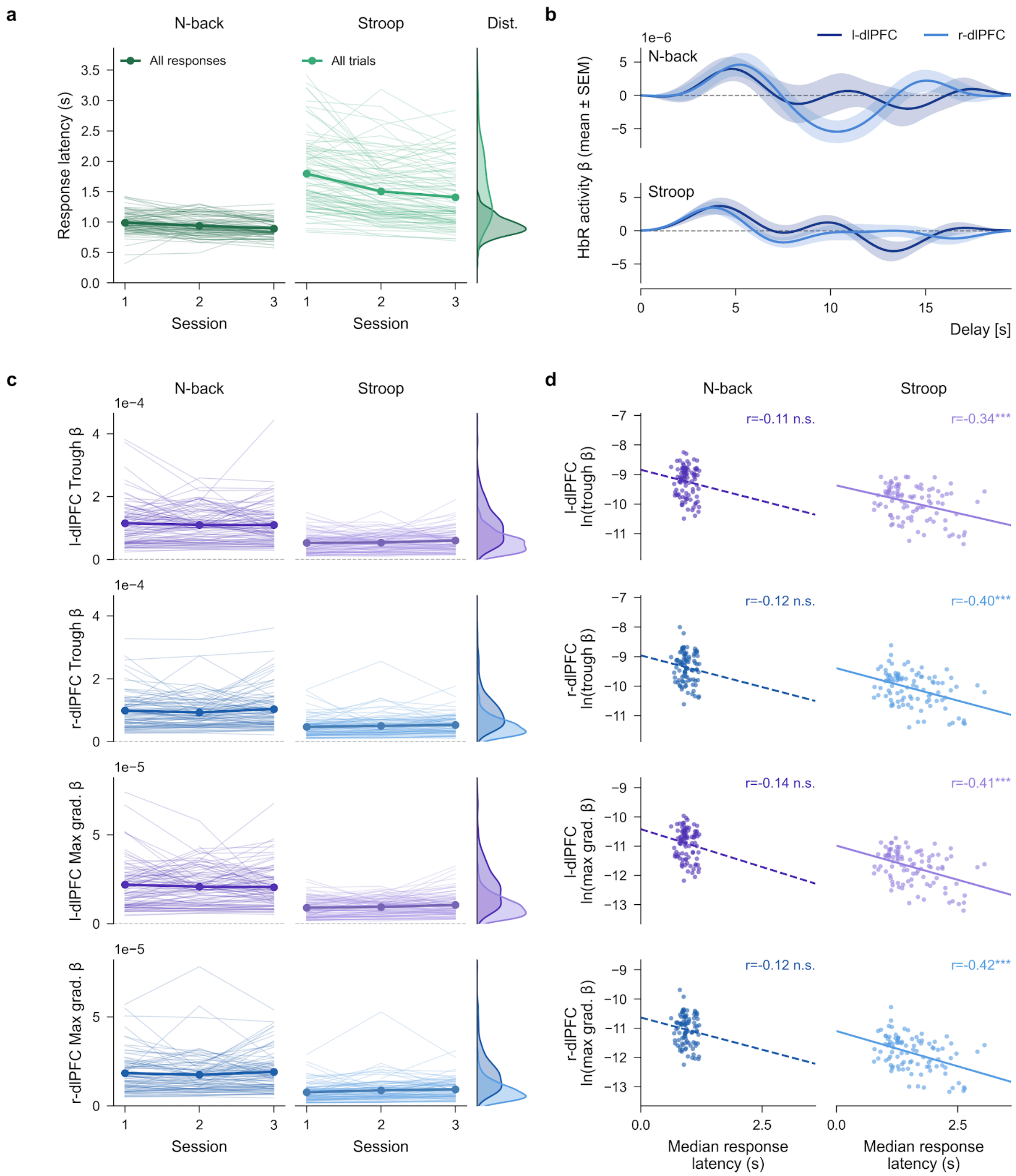


**Supplementary Figure 5.** HbR version of Figure 2 for PFC scaled data. (a) Response latency across sessions. (b) Mean HbR response. (c) Amplitude across sessions. (d) Amplitude against response latency.

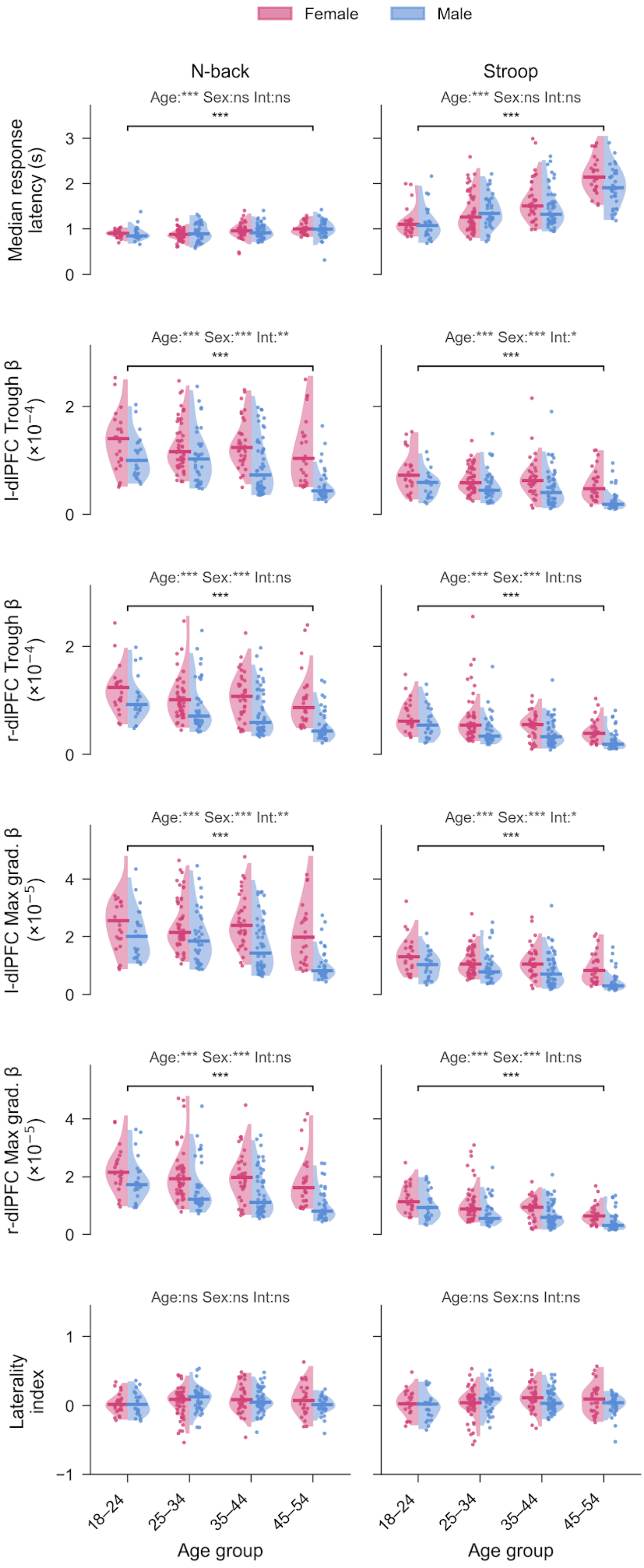


**Supplementary Figure 6.** HbR version of Figure 3 for PFC scaled data.
